# Narrative and Gaming Experience Interact to Affect Presence and Cybersickness in Virtual Reality

**DOI:** 10.1101/843169

**Authors:** Séamas Weech, Sophie Kenny, Markus Lenizky, Michael Barnett-Cowan

**Affiliations:** Department of Kinesiology, University of Waterloo, Waterloo, Ontario, Canada; The Games Institute, University of Waterloo, Waterloo, Ontario, Canada

**Keywords:** immersiveness, presence, cybersickness, motion sickness, virtual reality, user experience

## Abstract

Research has established a link between presence and cybersickness in virtual environments, but there is significant disagreement regarding the directionality of the relationship (positive or negative) between the two factors, and if the relationship is modulated by other top-down influences. Several studies have revealed a negative association between the factors, highlighting the prospect that manipulating one factor might affect the other. Here we examined if a top-down factor (narrative context) enhances presence, and whether this effect is associated with a decrease in cybersickness. We analyzed the association between responses to questionnaire measures of cybersickness and presence, as well as the degree to which their relationship was affected by the administration of an ‘enriched’ or ‘minimal’ verbal narrative context. The results of the first experiment, conducted in a controlled laboratory environment, revealed that enriched narrative was associated with increased presence, but that the reductive effect of narrative on cybersickness depended on video gaming experience. We also observed the expected negative association between presence and cybersickness, but only in the enriched narrative group. In a second experiment, conducted with a diverse sample at a public museum, we confirmed our previous finding that presence and cybersickness are negatively correlated, specifically when participants experienced an enriched narrative. We also confirmed the interaction between narrative and gaming experience with respect to cybersickness. These results highlight the complexity of the presence-cybersickness relationship, and confirm that both factors can be modulated in a beneficial manner for virtual reality users by means of top-down interventions.

## Introduction

Virtual reality (VR) applications have the capacity to make users feel as if they inhabit a world that is not their own. Although users who don a head mounted display are consciously aware that they will be a temporary visitor to an artificial space, users nonetheless engage with a simulation and respond to it in ways that resemble real-world reactions. A user can be startled when their virtual avatar is attacked (Lin, 2017), stressed when they are stalked by an artificial intelligence (Jin et al., 2016), awed when touring a virtual city (Guttentag, 2010), or nauseated when sitting in a simulated roller coaster (Davis et al., 2015). While VR is a powerful tool for evoking (and studying) natural behavior, the use of VR can also result in negative side effects.

Many VR users experience symptoms of physical discomfort that can include nausea, disorientation, and eye strain (LaViola Jr., 2000; McCauley & Sharkey, 1992). This experience is known as ‘cybersickness’ (Stanney et al., 1997). VR applications often introduce *sensory conflicts* (particularly visual-vestibular conflicts) which are thought by some to play a causal role in cybersickness (Oman, 1990; Reason, 1978; Rebenitsch & Owen, 2016; Weech et al., 2018a). Passive self-motion in VR is a key trigger for these conflicts, where the visual flow produced by viewpoint movement is unaccompanied by, or incongruent with, the expected vestibular and proprioceptive cues that typically emerge during self-motion. Equally, physical movement of the observer can be met with unexpected visual flow cues in some VR applications, with similar nauseogenic results (Palmisano et al., 2017; Stanney & Hash, 1998).

Some have proposed that the triggering of emesis by sensory conflict serves an adaptive function. It is possible that the nervous system attributes sensory conflict in VR to the ingestion of neurotoxic substances, and the emetic response serves to expel the perceived dangerous substance (Treisman, 1977). Alternatively, cybersickness may occur as part of a warning system that prevents maladaptive actions from occurring in environments where the normal sensorimotor control loops are disturbed (Takahashi et al., 1997). This theory proposes that the ‘retreating’ caused by cybersickness is adaptive as it prevents further harm to the individual.

The neurophysiological understanding of motion sickness has seen recent advances with the discovery of brainstem ‘sensory conflict neurons’ that respond primarily to unexpected passive motion (Brooks et al. 2015; Cullen 2014; Oman and Cullen 2014). Strong evidence exists that neural pathways involving vestibular nuclei underpins multiple forms of motion sickness (Balaban et al. 2014; Money & Wood, 1970; Yates et al. 1998, 2014). Cortical correlates of motion sickness include an increase in theta and delta band oscillatory activity in temporal and frontal brain regions (Chelen et al., 1993), particularly in the inferior frontal gyrus (Miller et al., 1996).

While the sensory conflict account of cybersickness posits a bottom-up cause for physical discomfort, the influence of top-down factors is also considered to be important. Experiential learning shapes expectations about sensory congruence, altering the impact of conflicting cues. For instance, long term exposure to a zero-gravity environment modulates the responses of astronauts to visual-vestibular mismatch: At first, a lack of a vestibular gravitoinertial force produces nausea, but this dissipates following a variable adjustment period (Reschke et al., 1998; Young et al., 1986). A similar habituation process is thought to occur with continued exposure to VR, where passive optic flow unaccompanied by vestibular situation will initially produce nausea, but symptom strength will reduce over multiple sessions (Stanney et al., 1999; van Emerick et al., 2011). Other explicit top-down influences have also been empirically linked to cybersickness. Particularly, presence, the sense of *being there* in the virtual world (Schuemie et al., 2001; Steuer, 1992; Sanchez-Vives & Slater, 2005), has been identified by many as a cognitive factor likely to modulate cybersickness.

While some groups have reported that the sense of presence in VR can increase cybersickness (e.g., Lin et al., 2002; Ling et al., 2013; Liu & Uang, 2011), we recently argued (Weech et al., 2019) that the balance of evidence suggests an inverse relationship between presence and cybersickness (e.g., Busscher et al., 2011; Cooper et al., 2016; Kim et al., 2005; Knight & Arns, 2006; Milleville-Pennel & Charron, 2014; Witmer & Singer, 1998). Several groups have claimed that this association results from a disruptive effect of cybersickness symptoms on presence, such that nausea and discomfort reorient attention away from a simulated environment (Bahit et al., 2016; Cobb et al., 1999; Nichols et al., 2000; Stróżak et al., 2018; Wilson et al., 1997; Witmer & Singer, 1998). At the heart of this theory is the idea that cybersickness diminishes presence by orienting attention inwards, away from the simulated environment.

Others have suggested that a strong sense of presence may direct attention away from the intrusive symptoms of cybersickness (Busscher et al., 2011; Cooper et al., 2016). The locus of attention has long been viewed as a central determinant of presence. Influential work by Witmer and Singer (1998) has proposed that presence requires an environment that captures the user’s attention, directing it away from external or internal cues that are unrelated to the simulated landscape. Factor analytic insights have suggested that attention may be one of two key cognitive processes required for VR presence, with the other process being the construction of a mental model to represent the simulated environment (Schubert et al., 2001; Vasconcelos-Raposo et al., 2016).

Experimental evidence suggests that manipulations of context can be used to achieve top-down effects on the sense of presence. For instance, presence is enhanced by the emotional context of a simulated environment (Baños et al., 2004). In a study by Baños and colleagues (2004), emotional content was provided in the form of ‘sad’ music and sentences (e.g., I have no future) that were heard as the participant explored a virtual park. The inclusion of high emotional context elevated feelings of presence and engagement compared to a neutral context. Although the authors found no significant difference between ‘neutral’ and ‘sad’ emotional context with respect to any negative side-effects (i.e., cybersickness), symptom severity was slightly lower in the ‘sad’ context. The authors highlight the psychological nature of presence, reiterating that physical immersiveness is but one requisite for achieving presence, and that top-down cognitive factors are also serious considerations (Baños et al., 2004).

Other research shows a significant impact of the perceived threat of a virtual environment on presence (Bouchard et al., 2008). By informing participants that an environment contained hidden snakes, Bouchard and co-workers (2008) produced increased levels of self-reported presence. The authors argued that a rapid, unconscious reaction to the phobic stimulus led participants to believe in the realism of the simulated world.

Smolentsev and colleagues (2017) recently documented evidence that presence is facilitated if a VR application begins with exposure to a ‘preamble’ environment that replicates the physical surrounding of the user. Congruence between the real and simulated environments was posited to result in a greater sense of continuity of memory and lower disconnect between the real and virtual environments.

It remains to be seen if a top-down intervention that increases presence also reduces cybersickness. This finding would be expected if one assumes that presence helps to distract from the negative side effects of sensory conflict in VR (Busscher et al., 2011; Cooper et al., 2016). On the other hand, if cybersickness were unaffected by a top-down manipulation to increase presence, this would indicate that the inverse presence-cybersickness correlation could be caused by the disruptive effects of sickness symptoms on presence.

Here we examined the effects of a top-down narrative manipulation on presence and cybersickness in a virtual environment. Participants underwent an audio narrative intervention prior to VR exposure that was either enriched, containing details about the nature of the simulated environment and the characters therein, or minimal, where the script was devoid of these details. Both narrative scripts contained closely matched details on the task, which was primarily to explore their space and interact with the characters. Following VR exposure, we measured levels of presence and cybersickness.

In addition, we were interested to examine the differences in susceptibility to narrative manipulation effects between populations who were experienced with video games, and those who were not. We predicted that individuals with high video gaming experience may be less affected by narrative manipulations due to a greater interest in identifying and exploiting the reward loop of the game, or simply due to lower initial levels of cybersickness than the non-gaming population. Furthermore, we aimed to partition the effects of narrative on the multidimensional experience of gameplay (termed ‘game engagement’; Brockmeyer et al., 2009) which is likely to be collinear with presence. To this end, we assessed the interaction between narrative and observers’ enjoyment, engagement, and sense of autonomy (altogether termed the ‘game experience’ of the user) following the VR exposure.

Rather than employing ‘explicit’ cues to modulate attention (e.g., “this object is important”), we expected the top-down narrative manipulation to generate ‘implicit’ cues that orient attention towards the virtual environment and away from the physical surrounding of the user (Nielsen et al., 2016). Several previous studies support the idea that implicit affective cues about an environment (e.g., whether an environment poses a threat) can modulate attentional selection, tuning perception without necessarily evoking conscious feelings (Friedman & Förster, 2010). In line with this evidence, we proposed that attention is likely to be shifted by an enriched narrative that produces a more compelling sense of presence, away from the effects of sensory conflict and towards the simulated environment. This attentional shift was expected to reduce the magnitude of cybersickness due to a decreased focus on internal symptoms of discomfort. Observing the expected pattern of results would support the idea that narrative manipulations pose a useful tool for protecting against cybersickness in virtual environments.

## Methods

### Experiment 1: Research Lab

#### Design

Testing was conducted at the Games Institute within the University of Waterloo. Participants were seated at a distance that allowed them to move their arms and lean forward without touching any objects in their physical surroundings (Figure 1). We used a commercial head mounted display (Rift CV1, Oculus VR; 90 Hz refresh rate, 1080 × 1200 resolution per eye) to present the Virtual Reality (VR) environment that was rendered by a computer with a high end graphics card (NVIDIA GTX1070). Inertial and optical sensors (Oculus VR sensors) tracked the orientation and position of the headset and controllers, and the head movement was used to update the viewpoint in VR. The capture space (~70 × 70 inches) and the inter-pupillary distance was calibrated for each participant using the packaged calibration software. The three motion capture sensors were placed around a circular chair allowing the participant to move 360° while seated and have their movements captured by the sensors.

**Figure 1.**
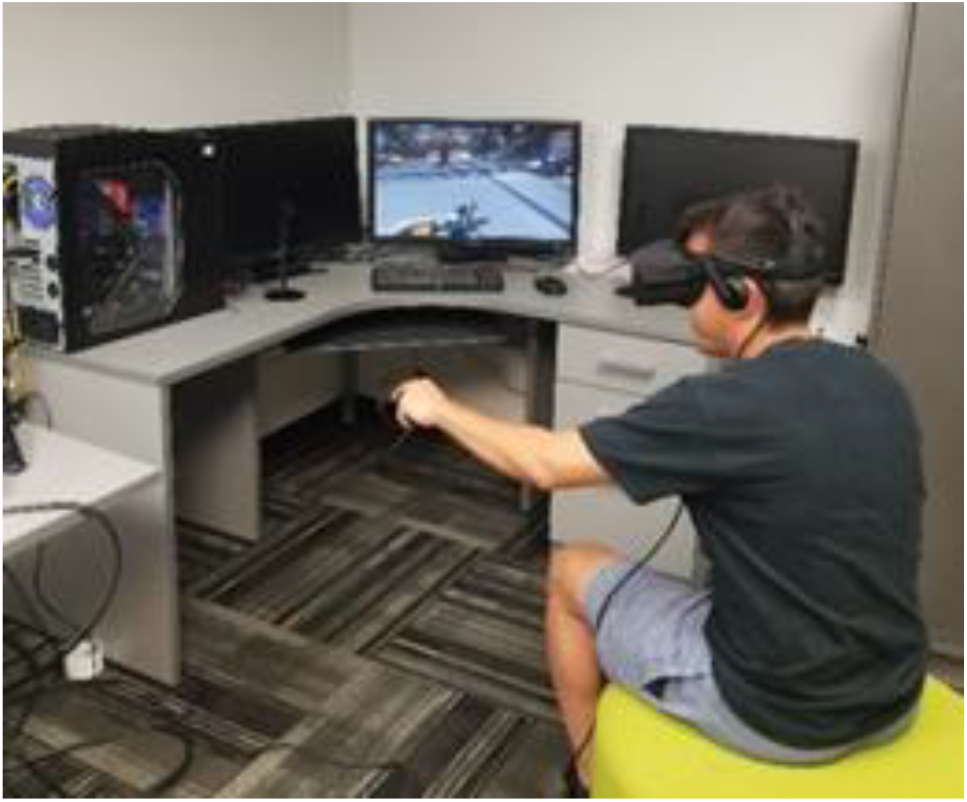
Depiction of a participant as they completed the VR portion of the experiment.

Participants filled out questionnaires on a table across from the VR set up before and after their gameplay experience. Participants were allocated into one of two conditions where different narrative contexts were provided prior to entering the virtual environment. Group assignment was random at first, and then assignment was based on previous video game experience in order to achieve equal group sizes.

##### Narrative Context

As a result of the group designation, participants were asked to listen to sound clips that provided either highly enriched narrative context (‘high’ condition), or a sound clip that provided minimal narrative context (‘low’ condition) as an introduction to the VR experience. This narrative context condition would provide the participant with either more (high) or less (low) background information with respect their impending experience. During the creation of the sound clips we ensured that the vocal tone and cadence were matched as well as possible. Due to the variable amount of information between the two contexts, the clip durations differed: 49 sec for enriched narrative context, compared to 28 sec for the minimal narrative context. Examples of the differences between the enriched and minimal narrative context conditions are presented in Table 1. Both sound clips are provided as Supplementary Material to this manuscript (available from the Open Science Framework: https://osf.io/kue7a/).

**Table 1.**
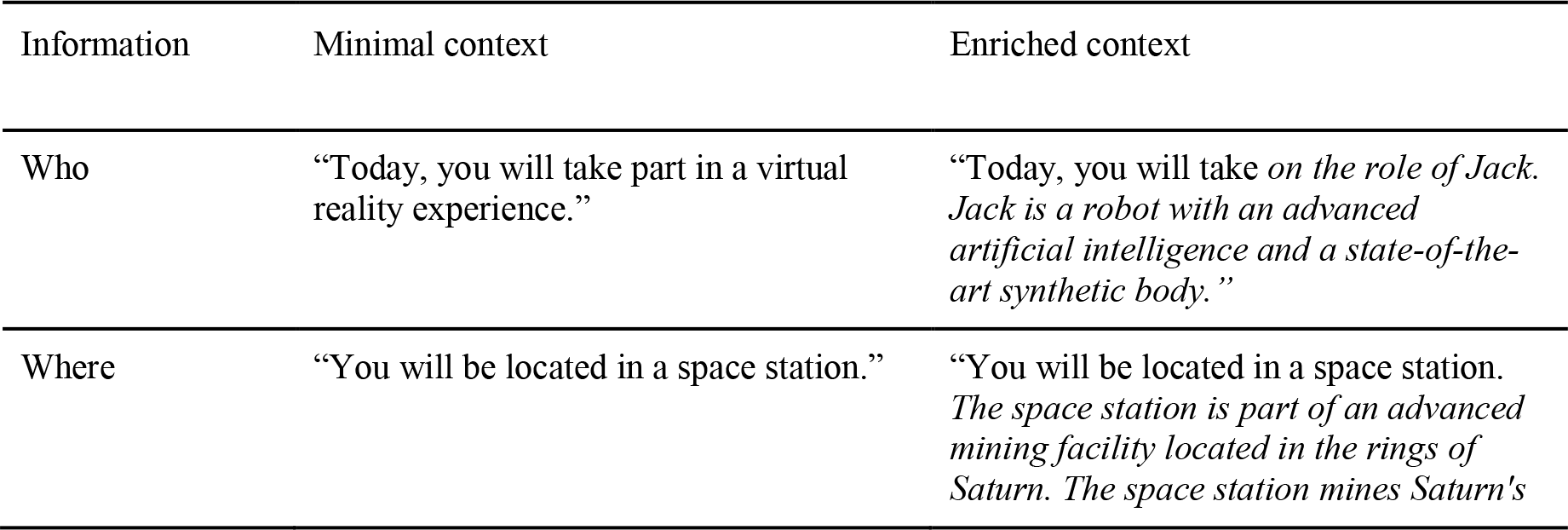
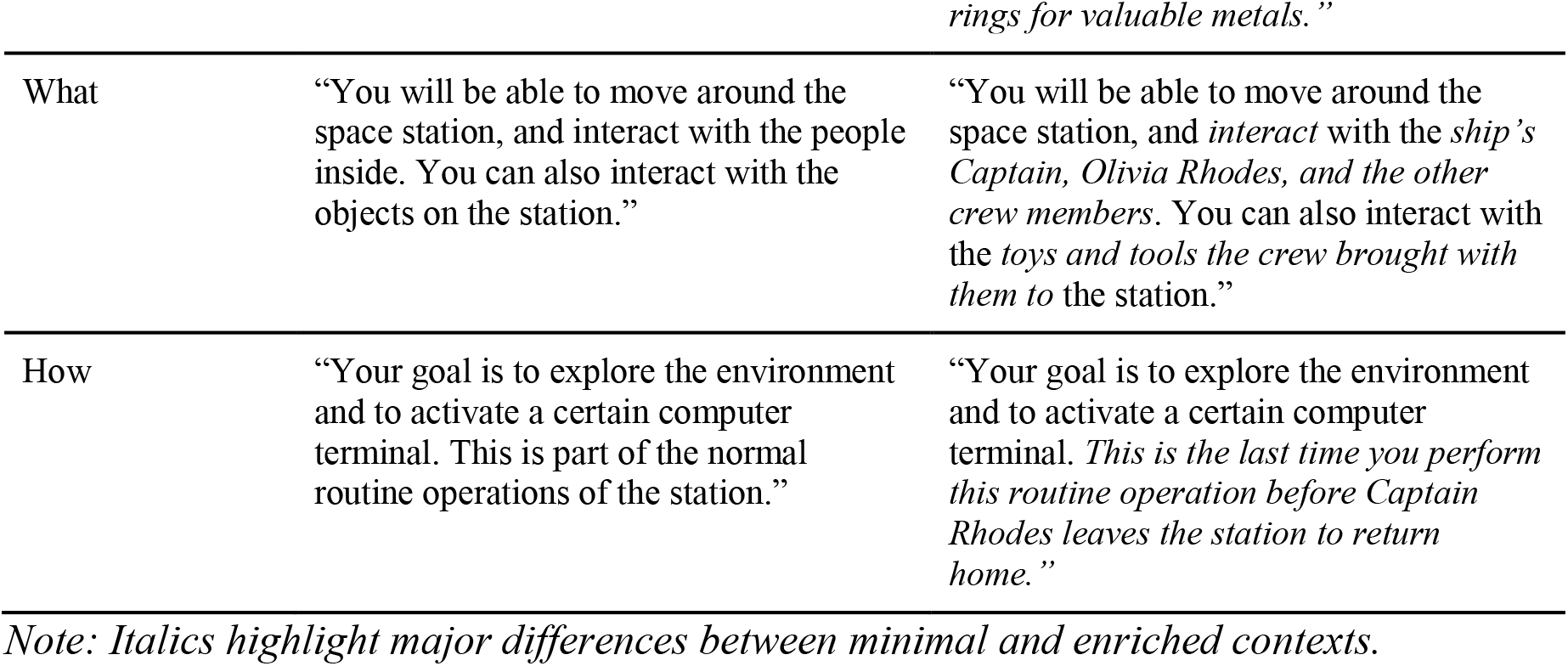
Details of Narrative Context Differences.

##### VR Experience

Participants were to play a section of the game Lone Echo (Figure 2A; Ready At Dawn, Oculus Studios). Lone Echo has received a ‘moderate’ comfort level rating on the Oculus store regarding its potential to induce cybersickness. Within the game, participants have a first-person view of a space station from the perspective of a robot. The participant controlled the robot by grabbing hold of surfaces and using push/pull actions to transport through the environment. Participants could explore the space station and interact with various objects and other human characters by using hand gestures to select dialog options. Movements and interactions were achieved through use of the hand-held controllers which were motion tracked by the sensors placed around the participant. The method of interaction was explained to participants before entering VR using an instructional video (Figure 2B).

**Figure 2.**
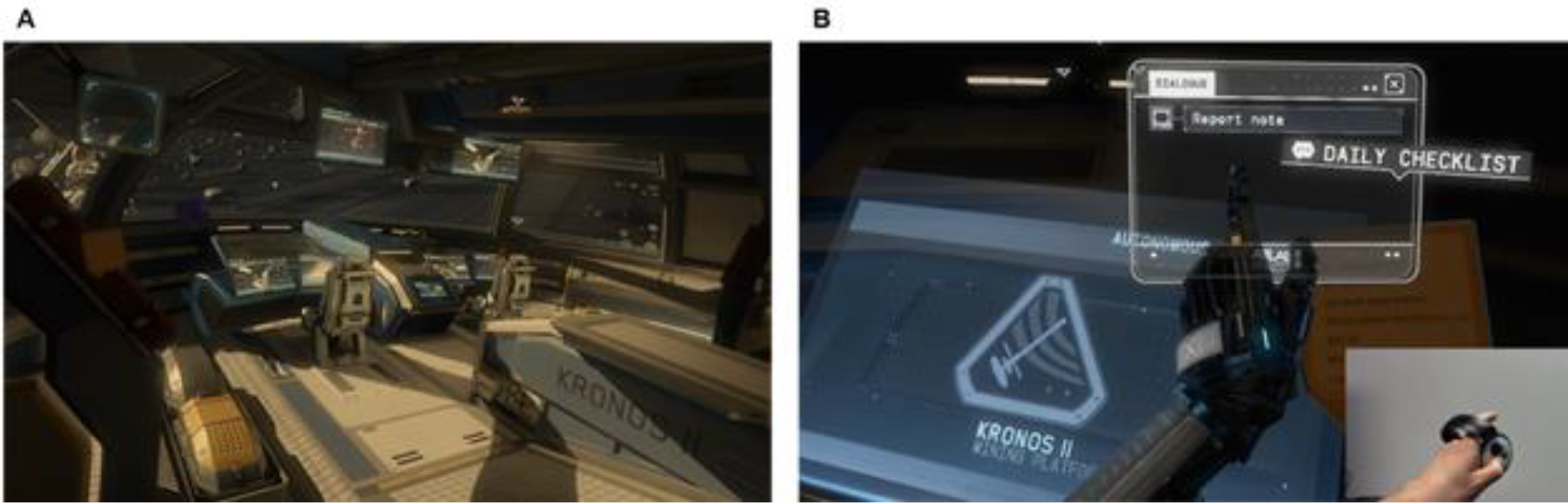
Screenshot of A) participants’ environment in the VR portion of the study, B) instructional video used to indicate controls to the participant.

#### Participants

42 volunteer participants (27 males and 15 females, ages were between 18 and 36, *M* age = 21.74, *SD* age = 3.50) took part in the experiment in a quiet laboratory setting. Participants recorded their gaming experience as part of the study; 12 participants had 0 hours of video game experience per week, 10 had <5 hours, 10 had 5–9, 4 had 10–14, and 6 had 15+. None of the participants reported any current or previous history of neurological disorders. All participants reported having vision that was normal or corrected to normal.

#### Procedure

The experiment procedure is shown in Figure 3. Participants first read an information letter containing details about the study in addition to signing a consent form. A questionnaire was used to collect information about participant demographics and measures of average weekly video game-play experience. Participants were asked to view an instructional video that outlined how to use the controllers for a game they would play. The video showed participants specifically how to hold the controllers, move in the game and interact with objects.

**Figure 3.**
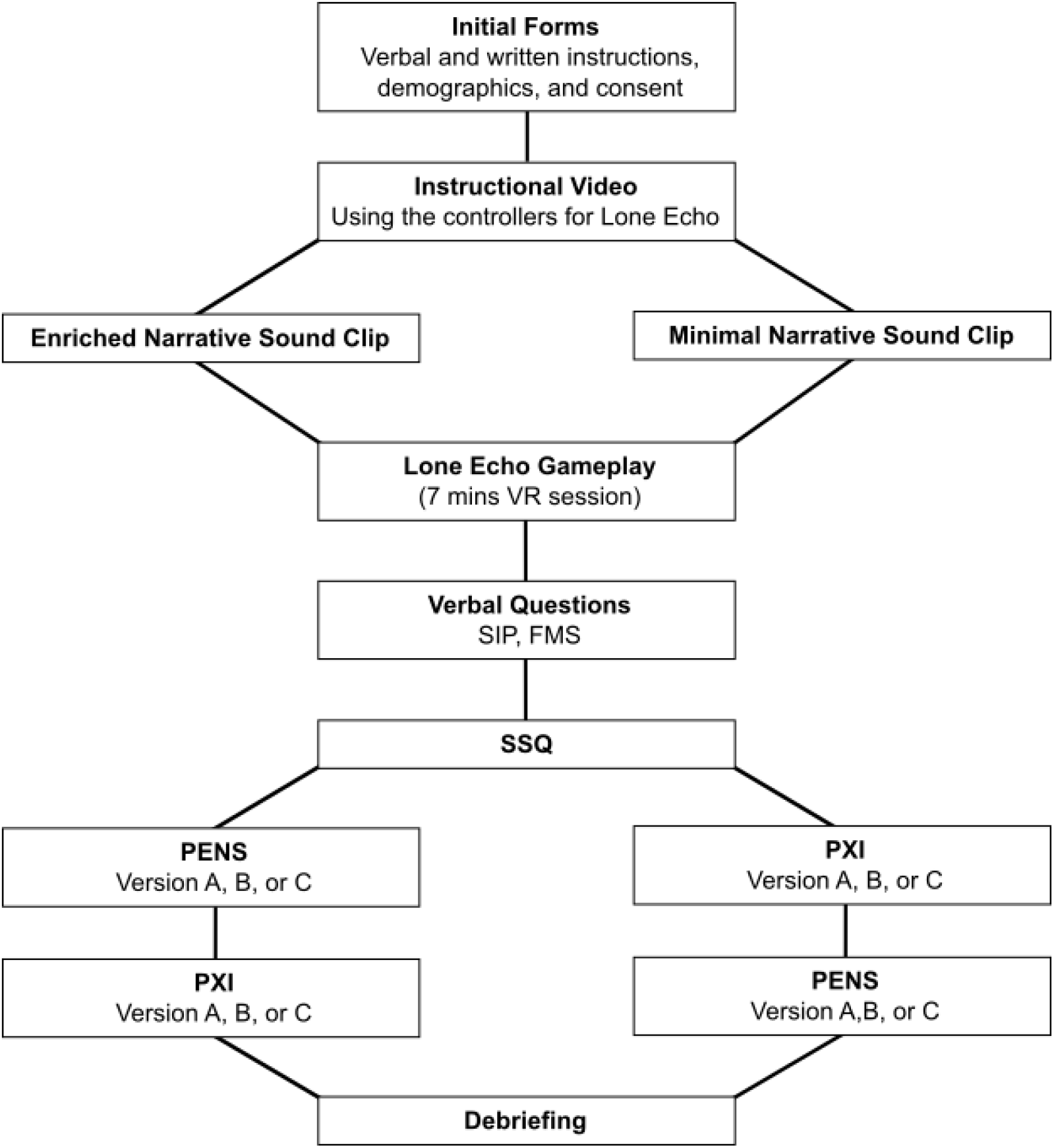
Schematic diagram of the experiment procedure.

Following calibration of the headset using the in-built software (head position and inter-pupillary distance), participants were asked to close their eyes while a sound clip was played which contained either an enriched or minimal context narrative introduction to the game. Immediately following the sound clip, participants were asked to open their eyes and begin the VR portion of the experiment. The participant then played 7 minutes of the game Lone Echo from a pre-selected load state (“Calibration Interrupted”). Progress in the game was measured in terms of the time taken by the participant to reach pre-determined checkpoints. Game-play was monitored by the investigator for unexpected events such as exiting to the pause menu or using controls that were not outlined in the instructional video.

After the 7 minute playing session was concluded, the investigator asked the participant two verbal questions about their experience. One question required participants to report their level of sickness (“On a scale of 0 – 20 where 0 is ‘none at all’ and 20 is ‘severe sickness’, how do you feel?” Fast Motion Sickness scale, FMS; Keshavarz & Hecht, 2011) and the other reporting their feeling of presence in the environment (“On a scale of 0 – 10 where 0 is ‘not at all present’ and 10 is ‘totally present’, to what extent did you feel present in the environment, as if you were really there?” Single Item Presence scale, SIP; Bouchard et al., 2004). Participants were then asked to complete written questionnaires, first completing the Simulator Sickness Questionnaire (SSQ; Kennedy et al., 1993) and then either the Player Experience of Need Satisfaction (PENS; Ryan et al., 2006) or another questionnaire, the Player Experience Inventory (PXI; Vanden Abeele et al., 2016). Note that answers on the PXI were used for the purposes of questionnaire validation and are not reported in the current experiment.

##### Post-Gameplay Questionnaires

After completion of the VR experience, the SSQ, PENS, and PXI questionnaires were administered to participants. The PENS and PXI each had three different versions which varied in question order. These different versions were assigned to participants randomly. The PXI and PENS were administered in a random order (SSQ was always first).

The SSQ involves rating 16 symptoms of sickness (e.g., ‘nausea’, ‘fatigue’) on a scale of ‘none’, ‘mild’, ‘moderate’, or ‘severe’ based on how the participant felt at the time. This questionnaire is a common measure for simulator sickness or cybersickness in terms of a total score and three sub-scores (nausea, oculomotor discomfort, and disorientation).

The PENS questionnaire consists of rating to what extent participants agreed with several statements about their experience playing the game within a 7-point scale ranging from ‘strongly disagree’ to ‘strongly agree’. The PENS questionnaire has been used to measure player satisfaction through assessing basic psychological needs related to competence, autonomy and relatedness that can be satisfied through game-play (Ryan et al., 2006). Each subscale of the PENS reflects individual underlying factors related to game experience: PENS Autonomy (PENS-A) indicates freedom of choice for executing actions in the game; PENS Relatedness (PENS-R) demonstrates connectedness to other in-game characters; PENS Intuitive Controls (PENS-IC) reflects the ease-of-controllability of movement and environmental interaction; PENS Competence (PENS-C) reflects the degree to which a player feels able to meet the challenge of the game; and PENS Presence-Immersion (PENS-PI), our main measure-of-interest from this scale, which indicates an individual’s sense of ‘being there’ in the simulated environment.

The PXI questionnaire, which we collected here for a separate investigation, consists of rating the user’s experience with playing a game on a 7-point scale ranging from ‘strongly disagree’ to ‘strongly agree’. Since this questionnaire has not yet been validated experimentally, the answers obtained from this scale are not reported here.

### Experiment 2 (Public Research)

Following Experiment 1, we aimed to obtain a larger, more diverse sample of participants in order to test the generalizability of the effects we observed. In Experiment 2 we conducted a replication of the experiment with a large convenience sample of participants. The sample of participants in Experiment 2 demonstrated a high degree of heterogeneity with respect to age, gameplay experience, and other factors that were relatively homogeneous in the sample of participants in Experiment 1.

#### Participants

##### Enrollment

A convenience sample of 153 visitors to TheMuseum (Kitchener, Ontario) enrolled in experiment during the 2018 Ontario Mid-Winter break (March 12-16). Participants enrolled in the 30 min experiment on a volunteer basis. Participants were only permitted to enroll if they were over 8 years of age (no upper age limit). Guests of the museum were informed that there was an opportunity to be involved in the experiment from the museum staff, but the staff did not take part in recruitment directly. Visitors were only engaged in a detailed discussion about the experiment if they came to the information booth. Enrollment in the experiment occurred at the information booth staffed by trained research assistants, who ensured informed consent in the study according to the Declaration of Helsinki. Participants were led to a small waiting area where they completed the consent forms and demographic questionnaires. Once a testing booth became available for use, the participant was led by an investigator to the booth, where they would complete the study. The research protocol was approved by the University of Waterloo Research Ethics Board.

##### Assignment

Each participant was tested individually and randomly assigned to one of the two Narration conditions according to a pre-generated random-assignment list. Prior to exclusion, participants were evenly distributed in each narrative condition, with 77 (50.3%) participants assigned to the enriched narrative condition, and 76 (49.7%) to the minimal narrative condition.

##### Excluded from analysis

Ten participants discontinued participation due to severe sickness (*n* = 2), falling off their chair (*n* = 1), being scared of heights (*n* = 1), or for other unspecified reasons (*n* = 6). Eight additional participants did not appropriately receive the experimental manipulation, due to either non-compliance with proper headset wear (*n* = 5; e.g., constantly readjusting or removal of the headset), or non-compliance with the instruction to play the game in silence (*n* = 3; e.g., requesting to talk with the research assistant or engaging in discussion with nearby family members). Additionally, data from one participant were removed because the participant learned of the expected outcome of the experimental hypothesis prior to taking part in the experiment (*n* = 1). Finally, six additional participants were excluded because they had declined to complete either or both of the SSQ and PENS questionnaires.

##### Analyzed participants

In total, data from 128 participants were included and analyzed in the current dataset. Participants were 46 females (82 males), 18.2 years of age on average (*SD* = 13.2 years). Given that the majority of participants consisted of parents or children, the distribution of ages was strongly bimodal. The average age of participants younger than 18 years old (for whom consent was provided by a parent/guardian) was 10.8 years old (*SD* = 2.20 years old), and the average age of participants older than 18 years old was 37.1 years old (*SD* = 10.5 years old). Participants also had indicated the frequency with which they played video games on a weekly basis, and the reported mode was between 0 and 5 hours a day.

Although slightly more participants from the minimal narrative condition were excluded, the assignment to the narrative conditions remains approximately equal, with 66 participants assigned to the enriched narrative condition (51.6%) and 62 (48.4%) participants assigned to the minimal narrative condition. According to a chi-square test of independence, the enriched and minimal narrative condition consisted of an equivalent number of males and females, *X*^2^(1, *N* = 128) = 0.669, *p* = .413, of younger and older participants, *X*^2^(1, *N* = 128) = 0.001, *p* = .980, and of regular gamers/non-gamers, *X*^2^(1, *N* = 128) = 0.574, *p* = .449. Additionally, the proportion of males and females was equivalent in the younger and in the older participant groups, *X*^2^(1, *N* = 128) = 0.000, *p* = 1. However, sex was not independent of regular gaming experience, *X*^2^(1, *N* = 128) = 14.8, *p* < .001, with males being more likely than females to play video games for more than five hours a week. Similarly, participants under the age of 18 were more likely to regularly play video games than older participants, *X*^2^(1, *N* = 128) = 13.7, *p* < .001.

#### Procedure

The procedure of Experiment 2 was largely similar to that of Experiment 1. The main difference between the experiments was that data from participants in Experiment 2 were collected at a public museum (“TheMuseum”, Kitchener, Ontario) over the course of a one-week period. The testing procedure was identical in terms of the participants’ experience in VR and the questionnaires administered. However, a few key differences between this experiment and Experiment 1 are described below.

#### Testing

The testing area in Experiment 2 comprised of three booths that were separated by curtains (Figure 4). These booths were 2 × 4 m in area and housed the same type of workstation, head mounted display, and motion sensors as in Experiment 1. Participants were also guided through the same testing procedure for data collection, starting with the instructional video and ending with administration of post-game-play questionnaires.

**Figure 4.**
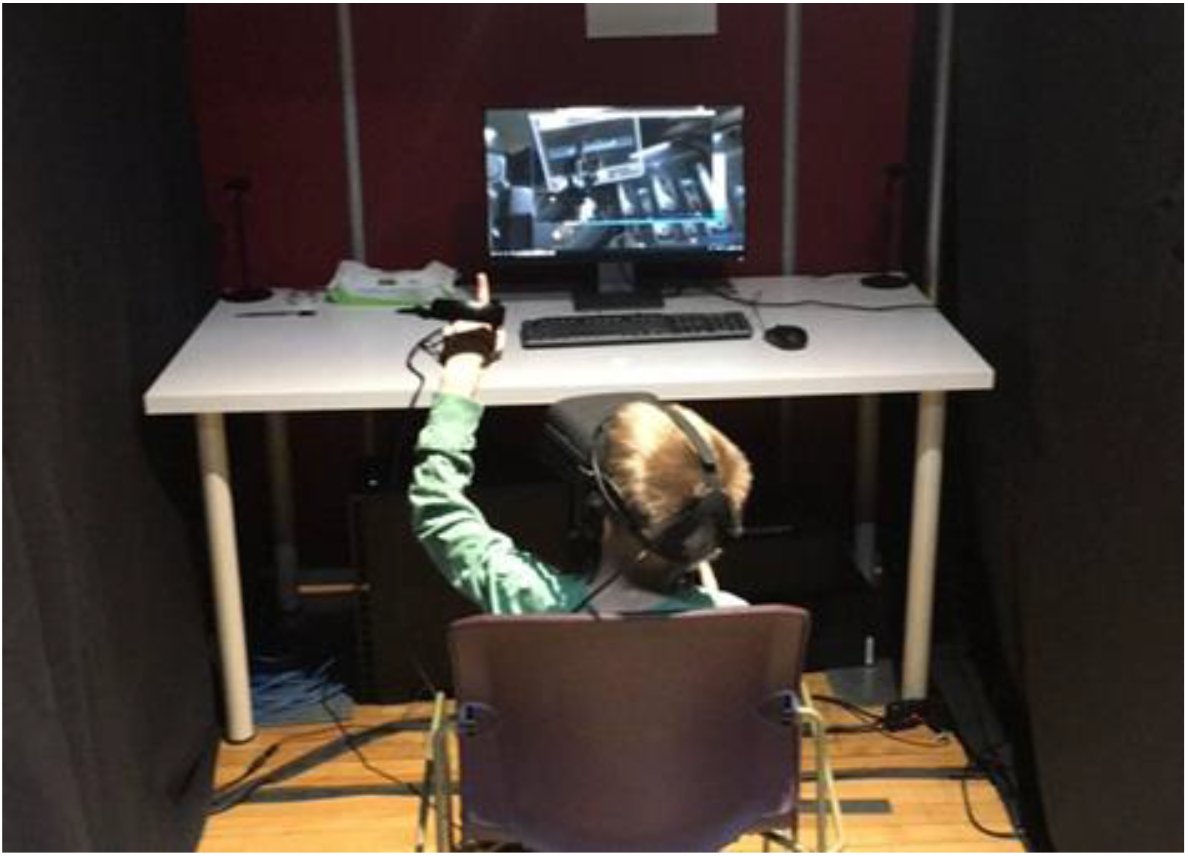
A participant experiencing the VR portion of the study in Experiment 2.

#### Post-Game-play Questionnaires

Questionnaires were completed in a quiet waiting area located beside the testing area. The procedure was similar to Experiment 1, but the SSQ was included in the random administration of questionnaires instead of always being completed first (as in Experiment 1). Otherwise, the randomization of the order of questionnaires remained the same. As in Experiment 1, study participation concluded once participants completed the final questionnaire and were provided with verbal and written debriefing.

## Results

### Experiment 1: Research Lab

#### A-priori analysis 1: There will be a negative correlation between presence (PENS-PI) and cybersickness (SSQ-T)

We first assessed if presence (PENS-PI) and cybersickness (SSQ-T) were correlated. As expected, we found a strong negative correlation between the two factors (Spearman’s ρ(40) = −.47, *p* = .002; Figure 5).

**Figure 5.**
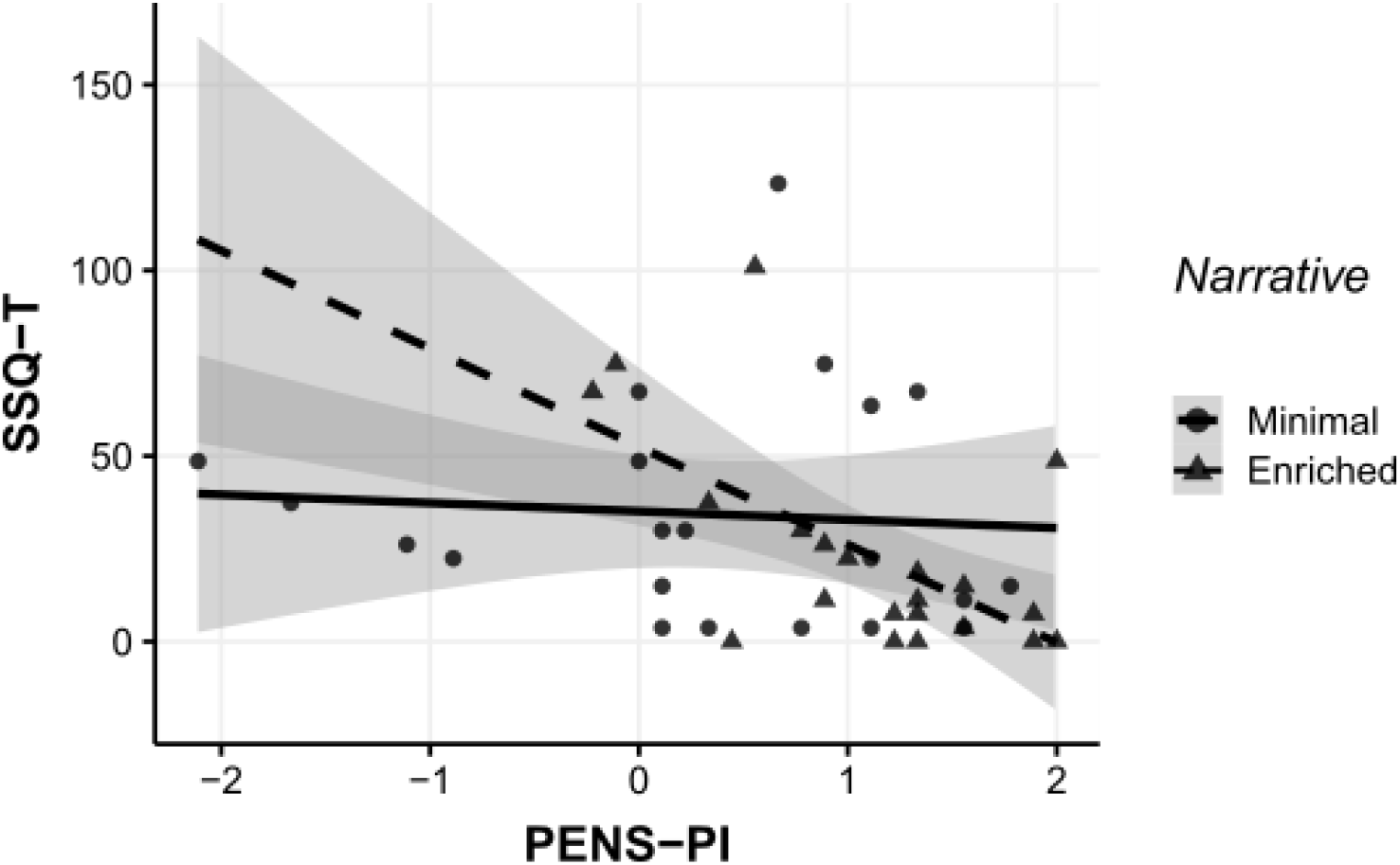
Scatterplot showing the relationship between presence (PENS-PI) and cybersickness (SSQ-T) measures across the minimal (circles, unbroken line) and enriched (triangles, dashed line) narrative groups in Experiment 1. Shaded area corresponds to 95% confidence intervals for the linear trends.

However, a post-hoc analysis revealed that when the data were separated by narrative condition, the correlation was only significant for the enriched narrative group (Spearman’s ρ(19) = −.51, *p* = .018) and not for the minimal narrative group (Spearman’s ρ(19) = −.25, *p* = .28).

#### A-priori analysis 2: Enriched narrative will result in lower cybersickness (SSQ-T) and higher in presence (PENS-PI) than minimal narrative

Given our prediction that a narrative intervention would result in increased presence and decreased cybersickness, we conducted independent samples Wilcoxon rank-sum tests with continuity corrections for the effect of narrative on PENS Presence-Immersion subscale scores (PENS-PI), and on SSQ Total scores (SSQ-T). We found that an enriched narrative led to significantly higher presence (*W* = 320, *p* = .013; Post-hoc analysis revealed power was .88). However, there was no significant difference in cybersickness between the enriched and minimal narrative groups (*W* = 161, *p* = .14; Post-hoc analysis revealed power was .32; Figure 6).

**Figure 6.**
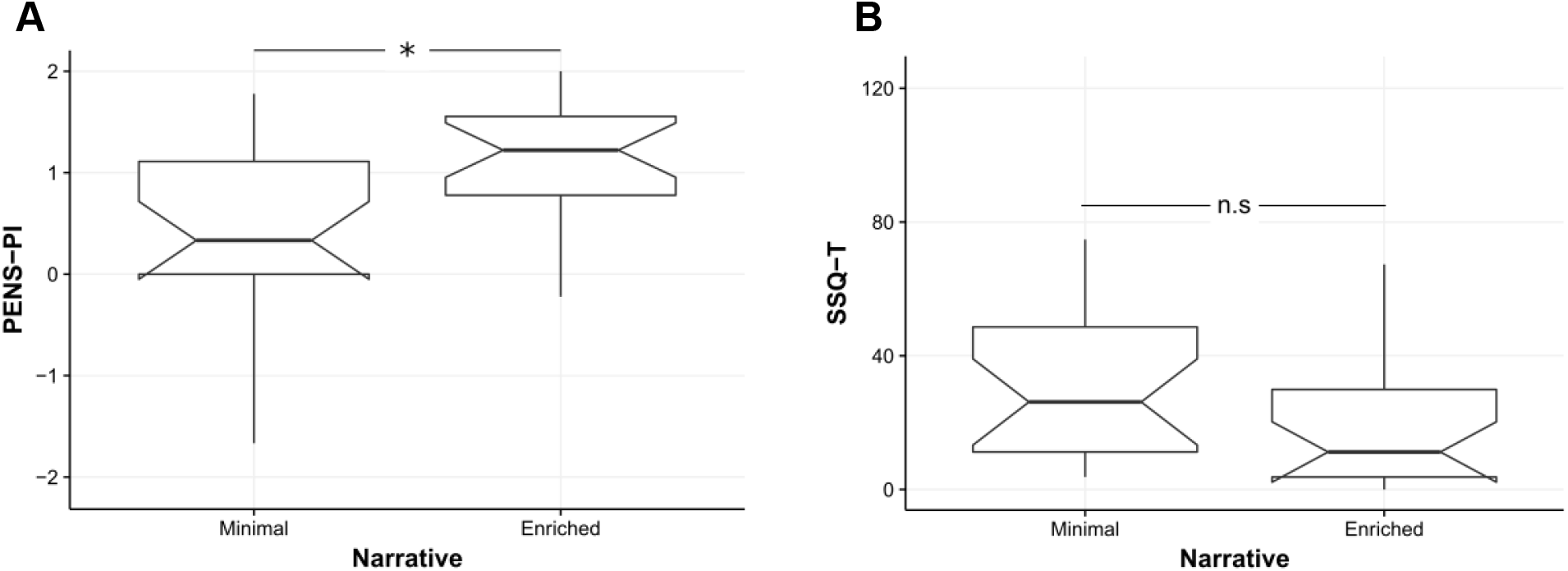
Notched Tukey boxplots indicating presence (PENS-PI, A) and cybersickness (SSQ-T, B) measures by narrative condition in Experiment 1. * *p* < .05.

#### Post-hoc analyses

We assessed relationships between measures of interest that were not specified a-priori. These include assessing the effect of narrative intervention on secondary outcome measures (i.e., measures other than PENS-PI and SSQ-T); correlations between other measures of cybersickness and presence; sex effects; and the effect of video game experience on our outcome measures (Figure 7).

**Figure 7.**
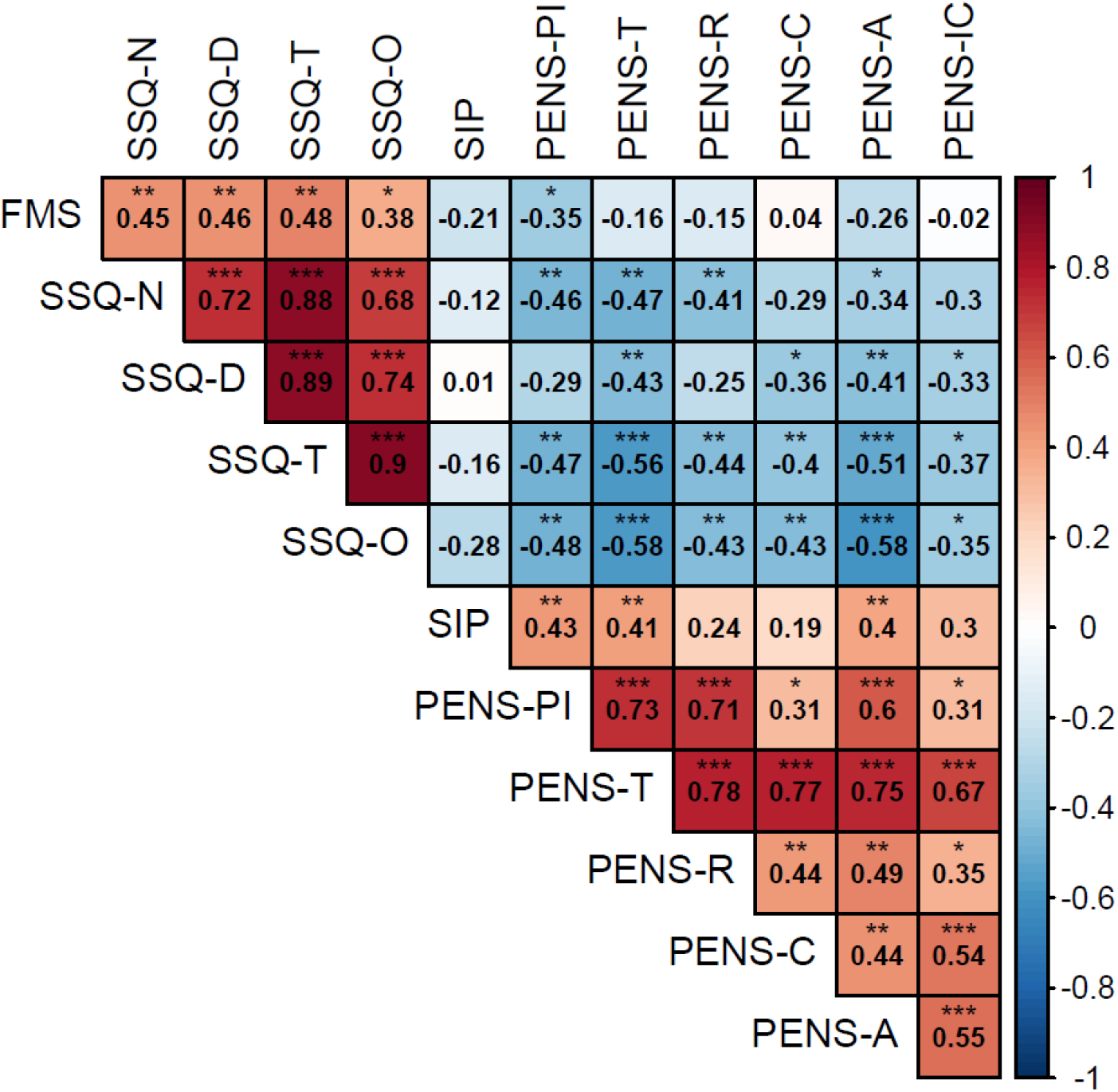
Hierarchically-clustered matrix of correlations (Spearman ρ(40) values) for outcome variables in Experiment 1. * *p* < .05, ** *p* < .01, *** *p* < .001.

##### Narrative manipulation

As well as the significant effect of narrative on presence discussed above, the enriched narrative intervention led to higher scores on the PENS-T (W = 310, *p* = .025; note that PENS-T encompasses the PENS-PI presence subscale, so these two effects are not orthogonal).

In addition to the null effect of narrative on SSQ-T described above, the narrative condition did not impact the following measures: SSQ-N (*W* = 283, *p* = .11) SSQ-O (*W* = 287, *p* = .09) SSQ-D (*W* = 247, *p* = .51), FMS (*W* = 197, *p* = .55), SIP (*W* = 270.5, *p* = .20), PENS-R (*W* = 295, *p* = .06), PENS-C (*W* = 296.5, *p* = .06), PENS-A (*W* = 254, *p* = .40), PENS-IC (*W* = 246.5, *p* = .52).

##### Goal Completion

Of the 42 participants, 10 successfully reached the specified goal during their VR exposure (i.e., they located and activated a computer terminal), while 32 did not. The narrative manipulation did not affect the likelihood that a participant would reach the terminal. 5 of 21 (24%) reached the terminal in both the enriched and minimal narrative groups (*X*^2^(1) = 0, *p* = 1).

Those who successfully activated the terminal reported higher feelings of competence (PENS-C: *W* = 73, *p* = .010), but also reported significantly higher disorientation (SSQ-D: *W* = 232.5, *p* = .030) and oculomotor discomfort (SSQ-O: *W* = 228, *p* = .041).

However, there were no differences between those who did/did not reach the terminal with respect to other cybersickness scores, presence, or game experience scores: SSQ-T (*W* = 224.5, *p* = .06), SSQ-N (*W* = 176.5, *p* = .63), FMS scores (*W* = 197.5, *p* = .26), SIP (*W* = 116.5, *p* = .19), PENS-T (*W* = 95.5, *p* = .06), PENS-R (*W* = 133, *p* = .43), PENS-IC (*W* = 123.5, *p* = .28), PENS-PI (*W* = 131, *p* = .40), or PENS-A scores (*W* = 103.5, *p* = .10).

##### Sex

Male participants scored higher on the intuitive controls subscale of the PENS than female participants (PENS-IC: W = 320, *p* = .002), and the PENS competence subscale (PENS-C: *W* = 313, *p* = .004).

Male and female participants did not differ with respect to the following measures: SSQ-T (*W* = 167, *p* = .36), SSQ-N (*W* = 179.5, *p* = .55), SSQ-O (*W* = 159, *p* = .25), SSQ-D (*W* = 167.5, *p* = .36), PENS-PI (*W* = 215, *p* = .75), PENS-R (*W* = 222, *p* = .61), PENS-A (*W* = 255, *p* = .17).

##### Additional measures of cybersickness and presence

We collected measures of cybersickness and presence in addition to the main outcome variables (SSQ-T and PENS-PI respectively). We compared the rapid measure of cybersickness (Fast Motion Sickness scale, FMS; Keshavarz & Hecht, 2011) with our main outcome measures, and found correlations with the SSQ-T (ρ(40) = .48, *p* = .001), SSQ-N (ρ(40) = .45, *p* = .003), SSQ-D (ρ(40) = .46, *p* = .002), and SSQ-O (ρ(40) = .38, *p* = .013).

With respect to the rapid measure of presence (Single Item Presence, SIP; Bouchard et al., 2004), we observed a significant positive correlation with PENS-PI scores (ρ(40) = .43, *p* = .005).

Although there was a negative correlation value for the association between the rapid SIP and FMS measures, we did not observe a significant relationship between the factors (ρ(40) = −.21, *p* = .18). A heatmap depiction reveals a large degree of clustering of responses at high presence and low cybersickness (Figure 8).

**Figure 8.**
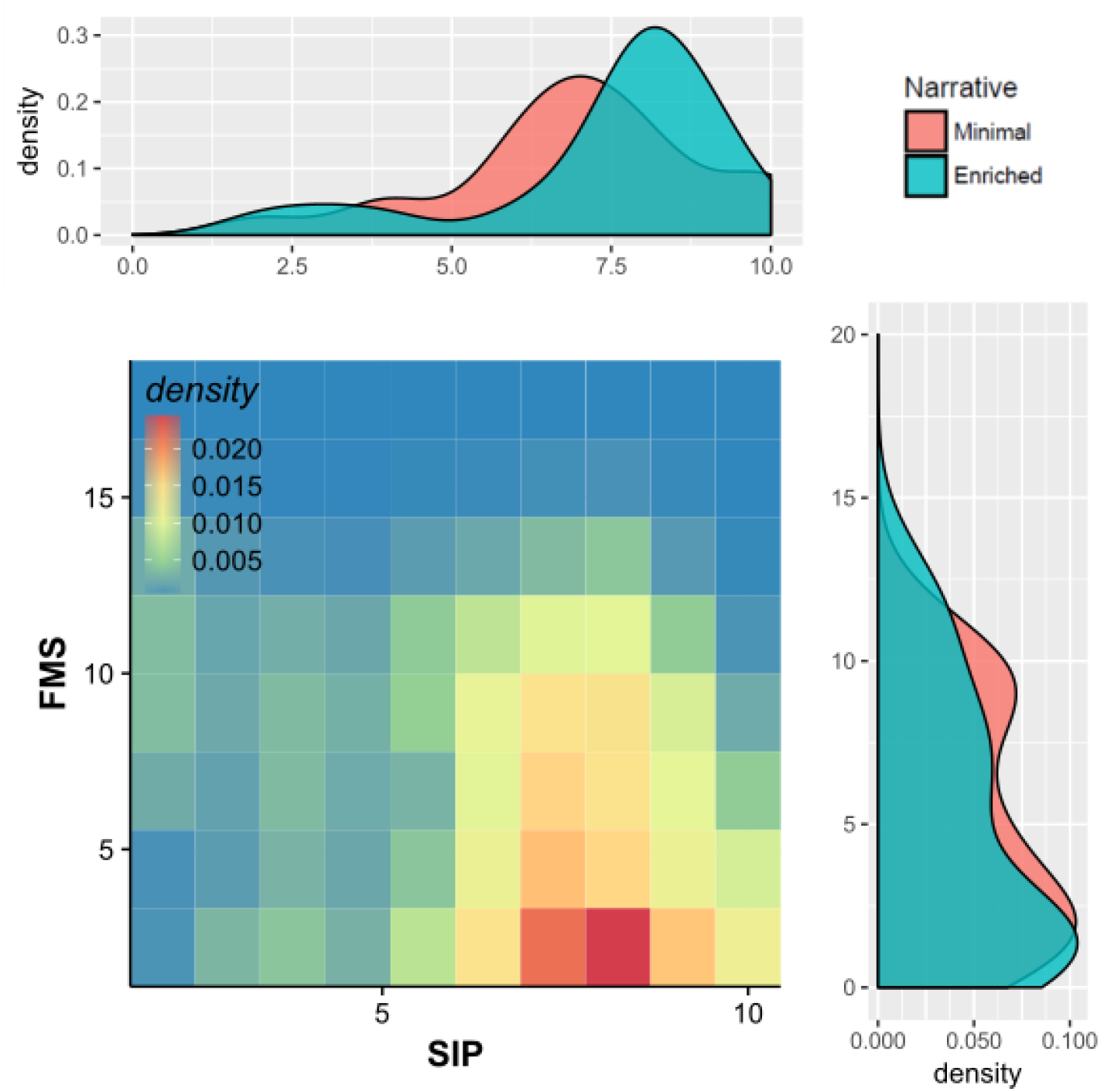
Heat-map (main) and density plots (top, right) for SIP and FMS responses in Experiment 1.

##### Regular Gamers vs. Non-Gamers

While there was no main effect of narrative on cybersickness, we observed a subtle but significant interaction between narrative and video game experience on SSQ-T scores (*F*(1, 38) = 4.15, *p* = .049; Figure 9). However, follow-up analyses revealed no difference in SSQ-T scores by narrative condition for any level of video game experience (*p*s > .17); as well, we found no significant difference between SSQ-T scores across levels of video game experience for the enriched (*F*(1, 19) = 1.07, *p* = .31) or minimal (*F*(1, 19) = 3.53, *p* = .08) narrative groups. We observed no correlation between video game experience and SSQ-T for the enriched (ρ(19) = .06, *p* = .79) or minimal (ρ(19) = −.36, *p* = .11) narrative groups. However, the number of participants in each condition was relatively low (e.g., of the 12 non-gamers, 8 were in the minimal group and 4 were in the enriched group).

**Figure 9.**
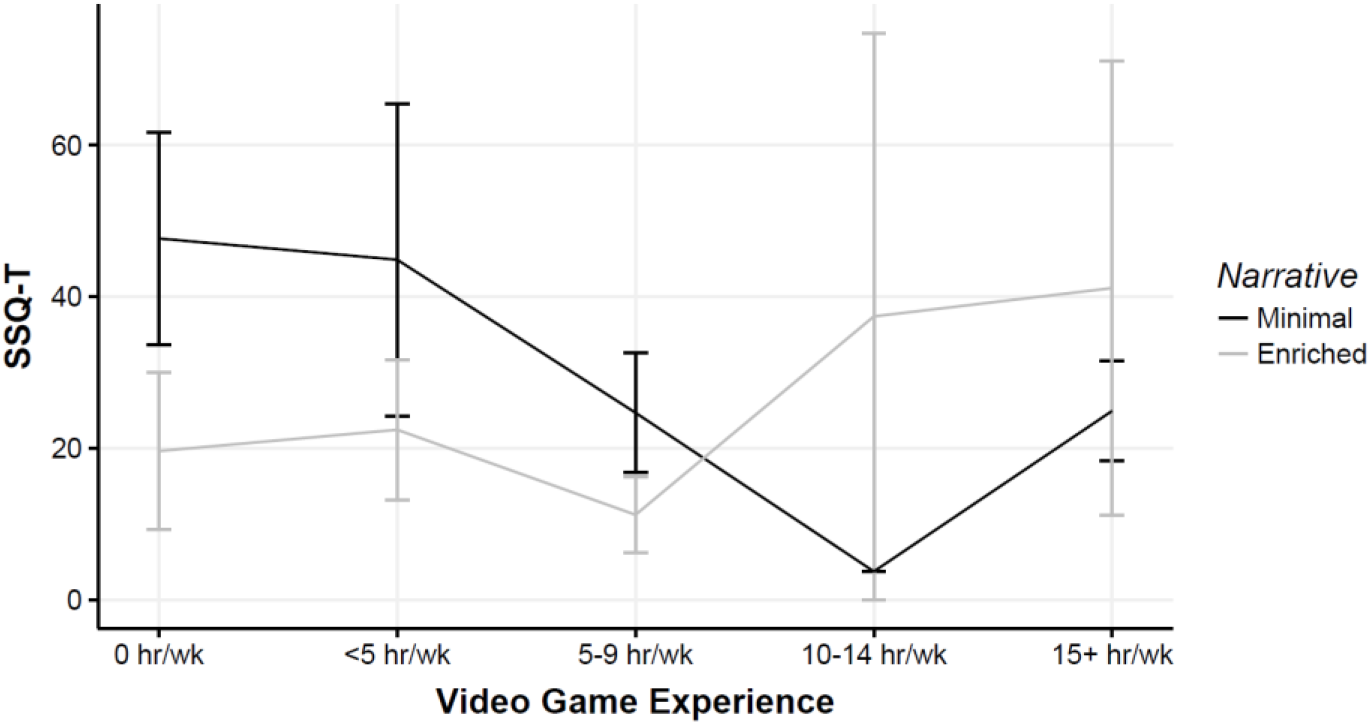
SSQ-T by video game experience and narrative group in Experiment 1.

##### Cybersickness Profile

Subscale scores for the SSQ are presented in Figure 10, with disorientation being the highest subscale score by mean and median. The second highest subscale was nausea, followed by oculomotor discomfort.

**Figure 10.**
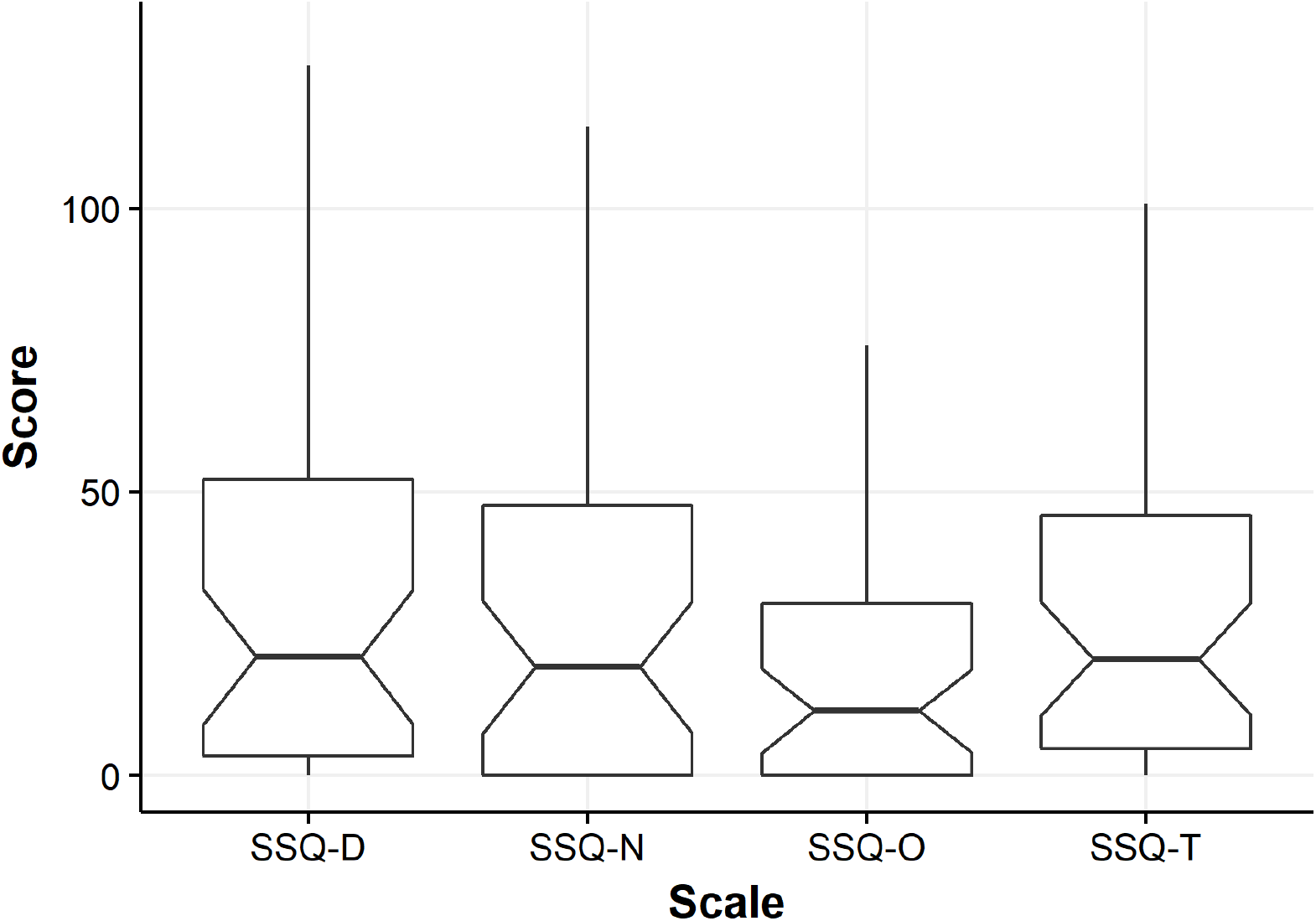
Notched Tukey boxplots showing cybersickness across subscales and total scores of the SSQ in Experiment 1. Bars are IQRs. D = Disorientation. N = Nausea. O = Oculomotor, T = Total.

### Experiment 2: Public Data Collection

#### A-priori analysis 1: There will be a negative correlation between presence (PENS-PI) and cybersickness (SSQ-T)

First, we tested if there was a correlation between presence (PENS-PI) and cybersickness (SSQ-T). In line with our predictions, and the results of Experiment 1, we obtained a significant negative correlation between the two factors (Spearman’s ρ(126) = −.26, *p* = .003; Figure 11).

**Figure 11.**
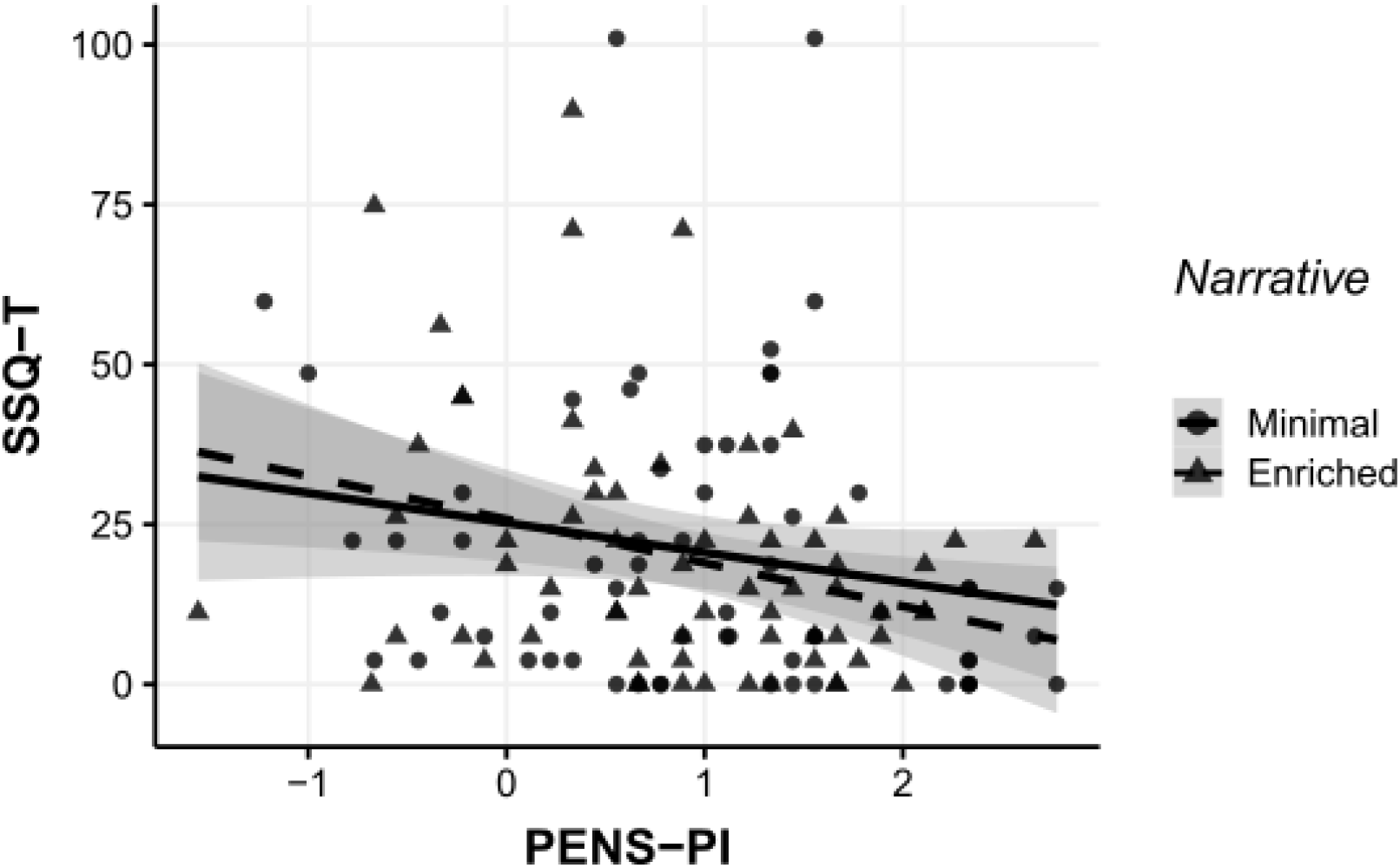
Scatterplot showing the relationship between presence (PENS-PI) and cybersickness (SSQ-T) measures across the minimal (circles, unbroken line) and enriched (triangles, dashed line) narrative groups in Experiment 2. Shaded area corresponds to 95% confidence intervals for the linear trends.

However, as in Experiment 1, when we analyzed data from the narrative groups separately, the correlation was only significant for the enriched narrative group (Spearman’s ρ(64) = −.28, *p* = .022) and not for the minimal narrative group (Spearman’s ρ(60) = −.24, *p* = .062).

#### A-priori analysis 2: Enriched narrative will result in lower cybersickness (SSQ-T) and higher in presence (PENS-PI) than minimal narrative

Contrary to our predictions, the narrative manipulation did not affect cybersickness (SSQ-T: *W* = 2118, *p* = .73; Post-hoc analysis revealed power was .07) or presence (PENS-PI: *W* = 1881, *p* = .43; Post-hoc analysis revealed power was .21; Figure 12).

**Figure 12.**
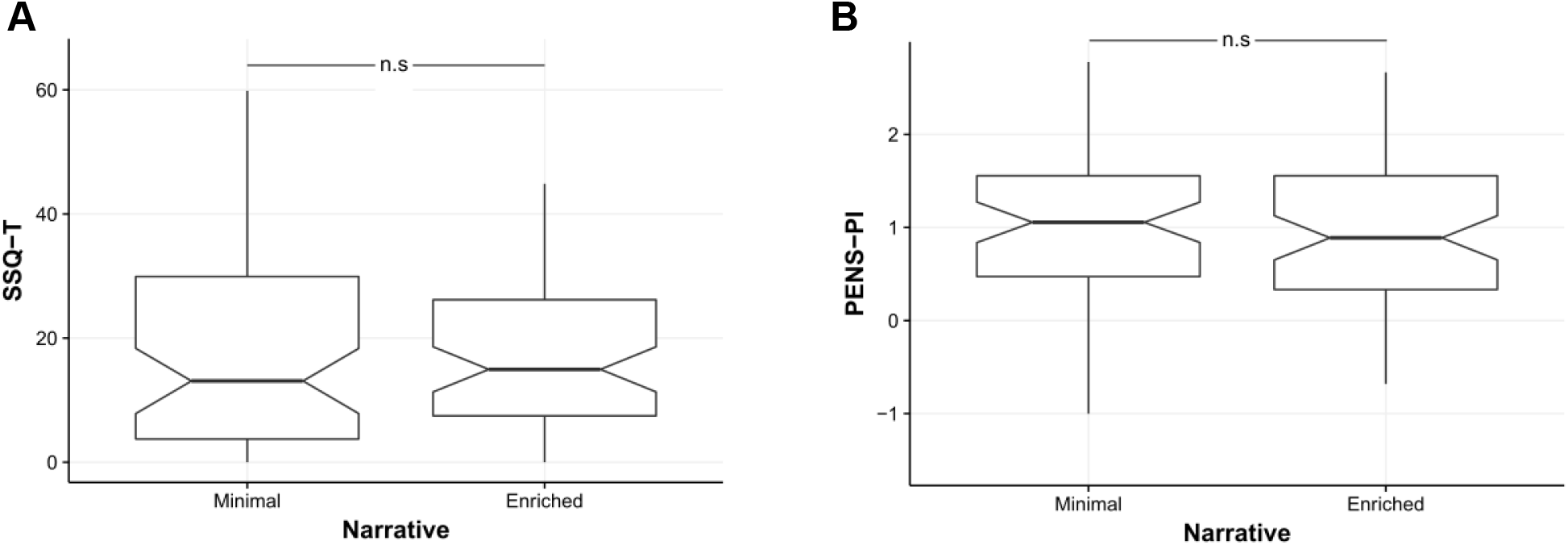
Notched Tukey boxplots indicating cybersickness (SSQ-T, A) and presence (PENS-PI, B) measures by narrative condition in Experiment 2.

#### Post-hoc analyses

As depicted in Figure 13, there were multiple significant correlations observed between outcome variables. In addition to the above analyses we conducted exploratory and post-hoc assessments of the effect of group and individual differences on multiple outcome variables that represented measures of presence, cybersickness, and game experience. The distributions of participants along these dimensions are depicted in Figure 14.

**Figure 13.**
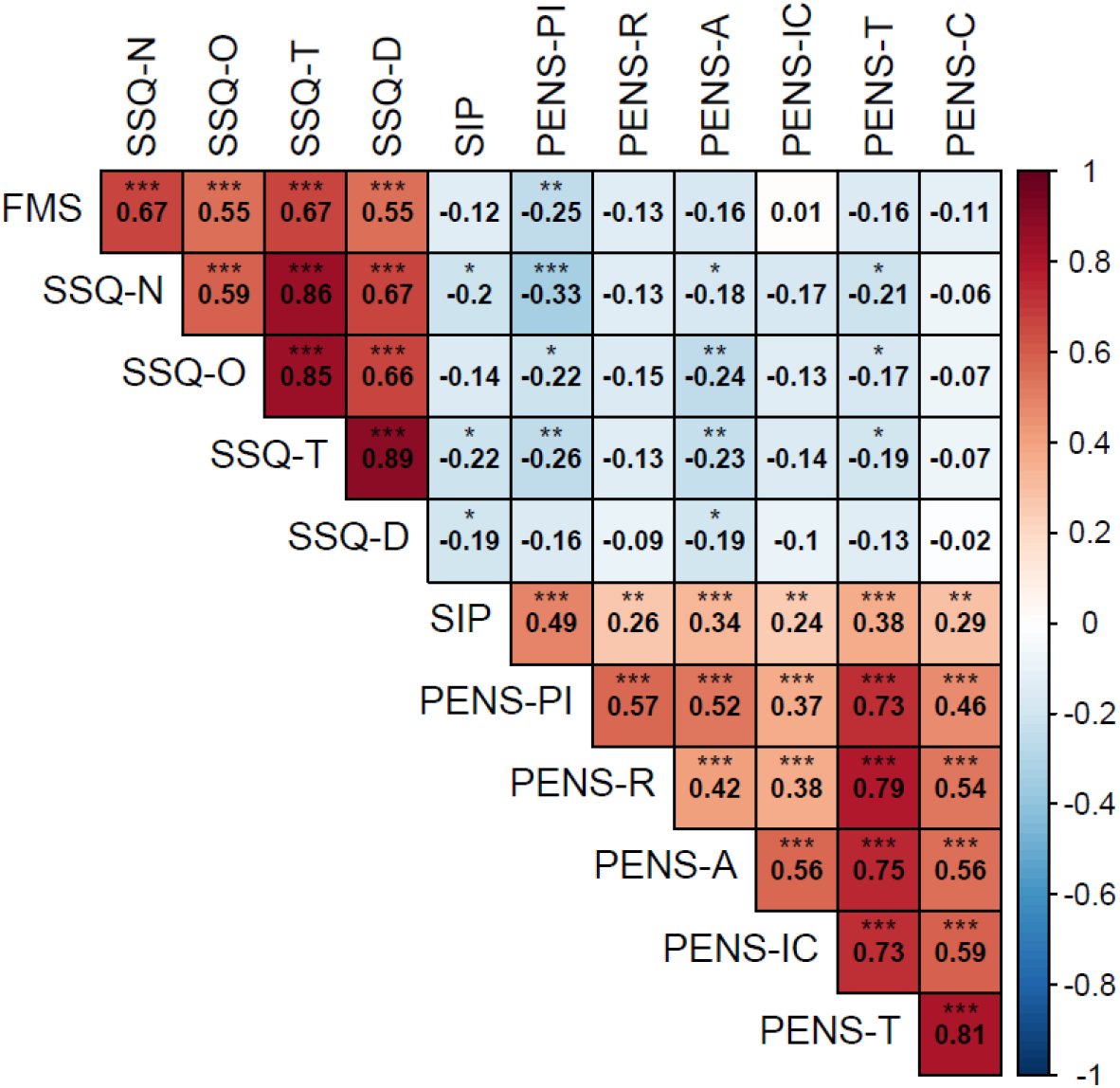
Hierarchically-clustered matrix of correlations (Spearman ρ(126) values) for outcome variables in Experiment 2. * *p* < .05, ** *p* < .01, *** *p* < .001.

**Figure 14.**
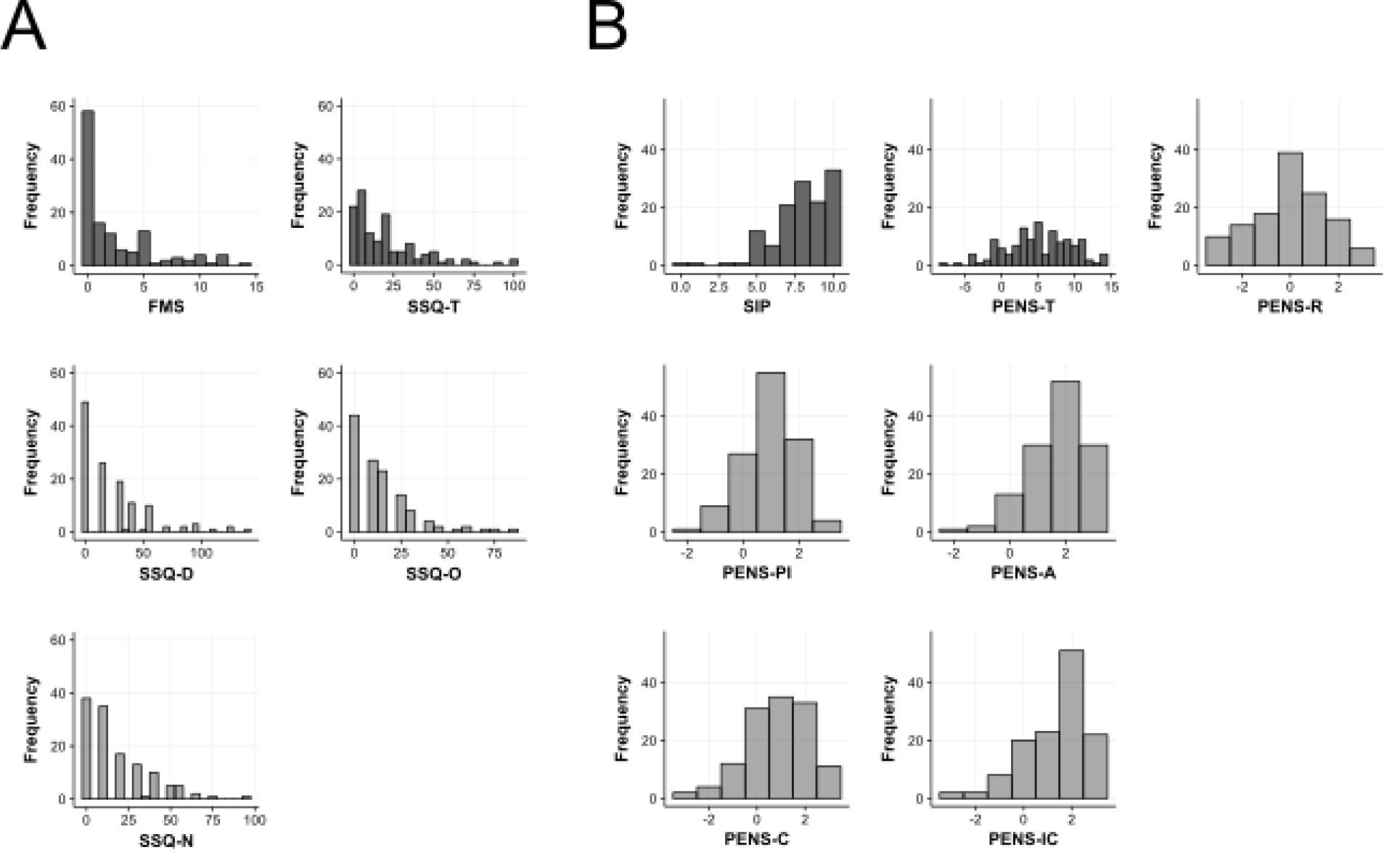
Histograms of responses in Experiment 2 with regard to A) Cybersickness; B) Presence and game-experience.

##### Narrative manipulation

In addition to the null main-effects of narrative context on the SSQ-T and PENS-PI, post-hoc analyses revealed no effect of narrative on any other measurements: SSQ-O (*W* = 2144.5, *p* = .63), SSQ-D (*W* = 2163, *p* = .56), SSQ-N (*W* = 2037.5, *p* = .97); FMS scores (*W* = 2283, *p* = .24); SIP scores (*W* = 1939, *p* = .60); PENS-T (*W* = 2125, *p* = .71); PENS-A (*W* = 2104.5, *p* = .78); PENS-IC (*W* = 2268.5, p = .29); PENS-C (*W* = 2179, *p* = .53); PENS-R (*W* = 2032, *p* = .95).

##### Additional measures of cybersickness and presence

We identified significant correlations between the FMS and SSQ-T (Spearman’s ρ(126) = 0.67, *p* < .001), as well as the subscales (SSQ-D: ρ(126) = .55, *p* < .001; (SSQ-N: ρ(126) = .67, *p* < .001; SSQ-O: ρ(126) = .55, *p* < .001). In addition, we observed a significant correlation between the SIP and the PENS-PI (ρ(126) = .49, *p* < .001).

The narrative manipulation did not affect these brief self-report measures (SIP: *W* = 2153, *p* = .60; FMS: *W* = 1809, *p* = .24).

With respect to the negative correlation between presence and cybersickness found with our main outcome variables, this relationship was not replicated for the FMS and SIP measures. As shown in Figure 15, the relationship between the FMS and SIP did not follow an inverse trend; rather, the measures both clustered highly at the extrema (low for FMS, high for SIP). Indeed, 15.6% of all participants rated themselves as 0 on the FMS scale and 10 on the SIP scale.

**Figure 15.**
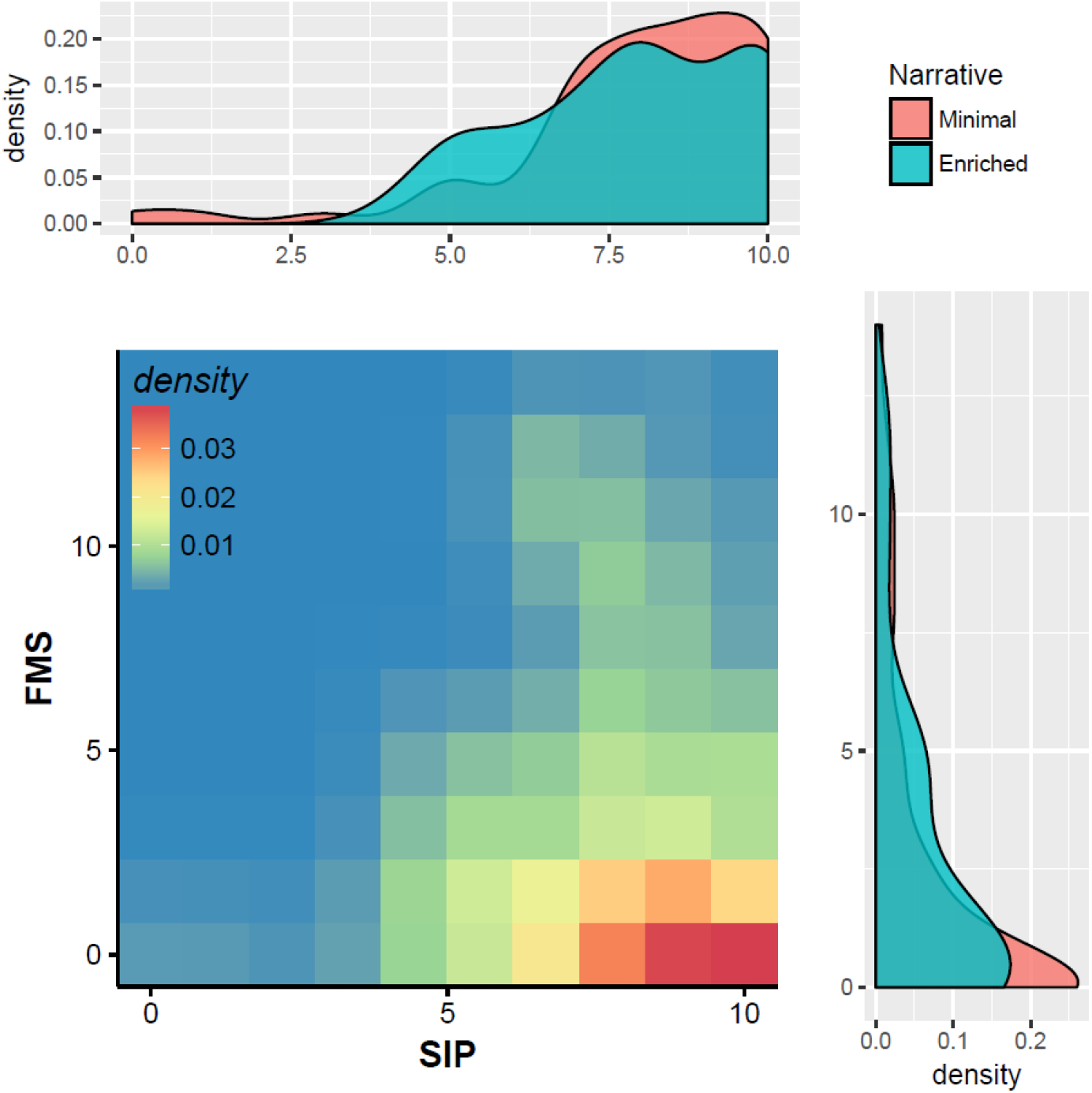
Heat-map (main) and density plots (top, right) for SIP and FMS responses in Experiment 2.

##### Regular Gamers vs. Non-Gamers

We characterized regular gamers as individuals who reported greater than 5 hours of video game play each week. In our sample, 70 were classified as gamers, while 58 were non-gamers. Gamers experienced less nausea and oculomotor discomfort (SSQ-N: *W* = 2464.5, *p* = .033; SSQ-O: *W* = 2499.5, *p* = .021), reported a greater intuitiveness of controls in the game (PENS-IC: *W* = 1521.5, *p* = .014), and reported higher competence (PENS-C: *W* = 1542, *p* = .019) than non-gamers.

However, the groups did not differ on any other outcomes: SSQ-T (*W* = 2394, *p* = .080), SSQ-D (*W* = 2122.5, *p* = .65), FMS scores (*W* = 2184.5, *p* = .44), SIP scores (*W* = 2251.5, *p* = .28), PENS-T (marginal; *W* = 1624, *p* = .052), PENS-R (*W* = 1652, *p* = .068), PENS-PI (*W* = 1998, *p* = .88), PENS-A (*W* = 1840, *p* = .36).

Importantly, there was a significant interaction between narrative and video game experience with respect to SSQ-T scores (*F*(4, 118) = 2.59, *p* = .04; Figure 16). A follow-up analysis showed that SSQ-T scores did not differ by narrative condition for any level of video game experience (*p*s > .11). At the same time, SSQ-T differed across the varying levels of video game experience, but only for the minimal narrative group (*F*(4, 57) = 4.00, *p* = .006). Specifically in this group, lower video game experience was associated with greater SSQ-T scores than higher video game experience (Spearman’s ρ(60) = −.33, *p* = .009). For the enriched narrative group, SSQ-T scores did not differ across varying levels of video game experience (*F*(4, 61) = 0.14, *p* = .97). We also saw no correlation between video game experience and SSQ-T for the enriched narrative group (ρ(64) = .03, *p* = .83). A caveat to these results involves the low number of participants in each condition (i.e., there were 18 non-gamers overall; only 8 non-gamer participants were in the enriched narrative group, while 10 non-gamer participants were in the minimal narrative group).

**Figure 16.**
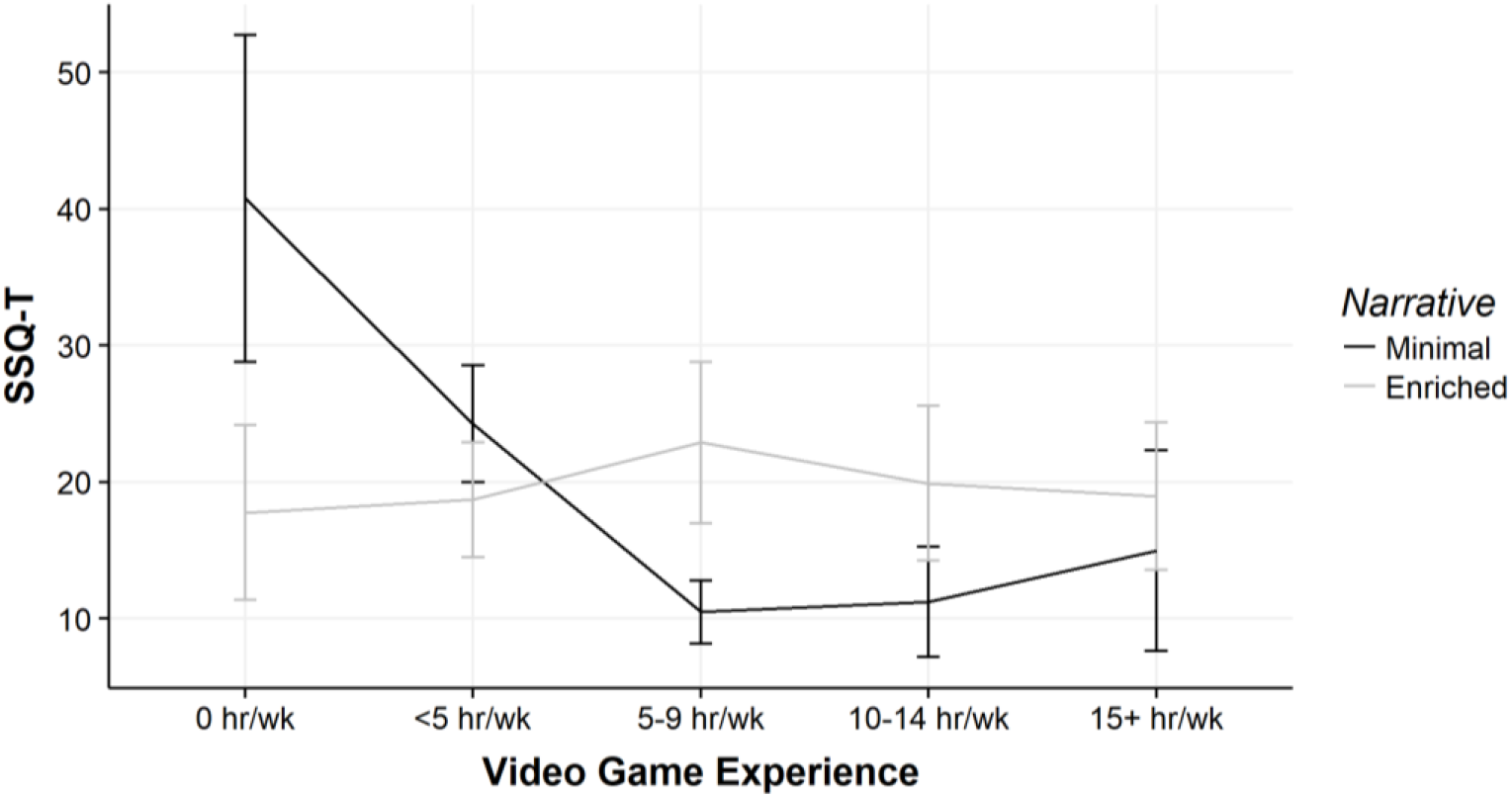
SSQ-T by video game experience and narrative group.

##### Goal Completion

Of the 128 participants, 76 successfully reached the specified goal during their VR exposure (i.e., they located and activated a computer terminal), while 52 did not. As in Experiment 1, the narrative manipulation did not affect the likelihood that a participant would reach the terminal. 41 of 62 (66%) in the minimal narrative group reached the terminal, whereas 35 of 66 (53%) in the enriched narrative group reached the terminal. This difference was not significant (*X*^2^(1) = 1.76, *p* = .18).

Those who activated the terminal scored significantly higher on PENS-T (*W* = 1478, *p* = .016), PENS-R (*W* = 1555, *p* = .039), and PENS-C (*W* = 1287, *p* < .001). However, there were no differences between those who did/did not reach the terminal with respect to other PENS scales (PENS-IC: marginal, *W* = 1576, *p* = .051; PENS-PI: *W* = 2066.5, *p* = .66; PENS-A: *W* = 1751.5, *p* = .27), or other measures of presence (SIP: *W* = 2270, *p* = .15) and cybersickness (SSQ-T, *W* = 1960, *p* = .94; SSQ-D, *W* = 1909, *p* = .74; SSQ-O, *W* = 1988, *p* = .95; SSQ-N, *W* = 1999, *p* = .91; FMS: *W* = 1778.5, *p* = .31).

##### Age

In our sample there were 92 participants under the age of 18 years old, and 36 over 18s (Figure 17).

**Figure 17.**
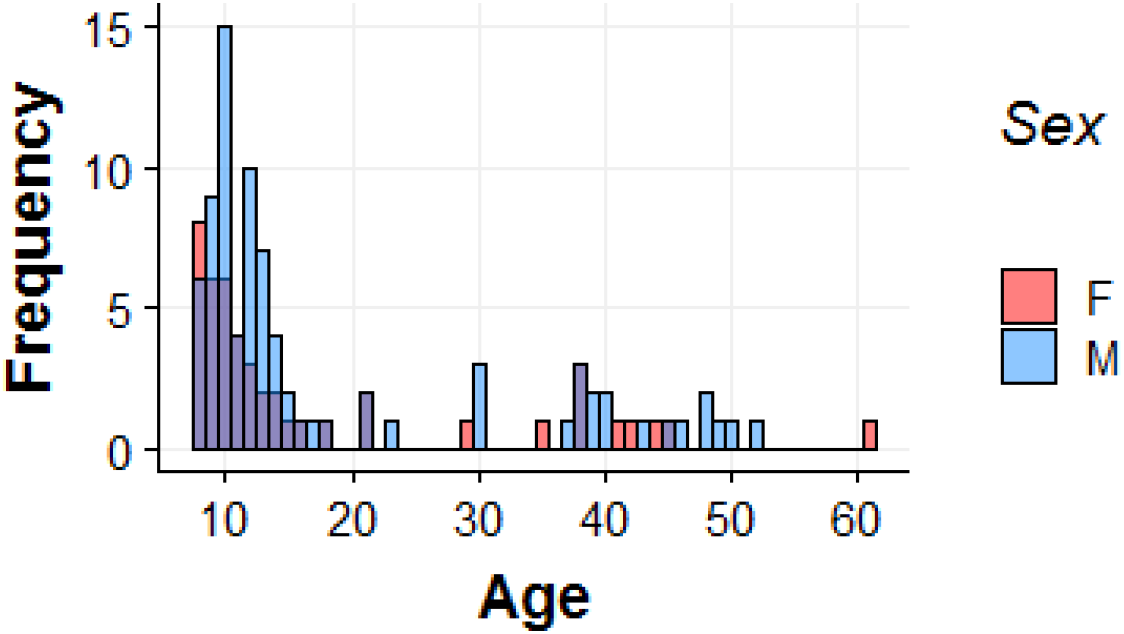
Histogram of ages (years) by participant sex (Red = Female; Blue = Male; Purple depicts overlap).

Given the possibility that the narrative context could have been more poorly attended by younger children than adults, we assessed the interaction between age and narrative context on the two main outcome measures. We found no evidence supporting an interaction between age and narrative context for the PENS-PI (*F*(1, 124) = 0.25, *p* = .62) or SSQ-T (*F*(1, 124) = 0.55, *p* = .46).

Under 18s scored higher on the PENS-T (*M* = 5.87, *SEM* = 0.46) than over 18s (*M* = 2.77, *SEM* = 0.76), resulting in significant group differences (*W* = 2248.5, *p* = .002) and a negative correlation between age and PENS-T (Spearman’s ρ(126) = −.23, *p* = .010; Figure 18). This relationship was underlied by significant negative correlations between age and the PENS-IC subscale (ρ(126) = −0.29, *p* < .001), as well as the PENS-C subscale (ρ(126) = −0.21, *p* = .017).

**Figure 18.**
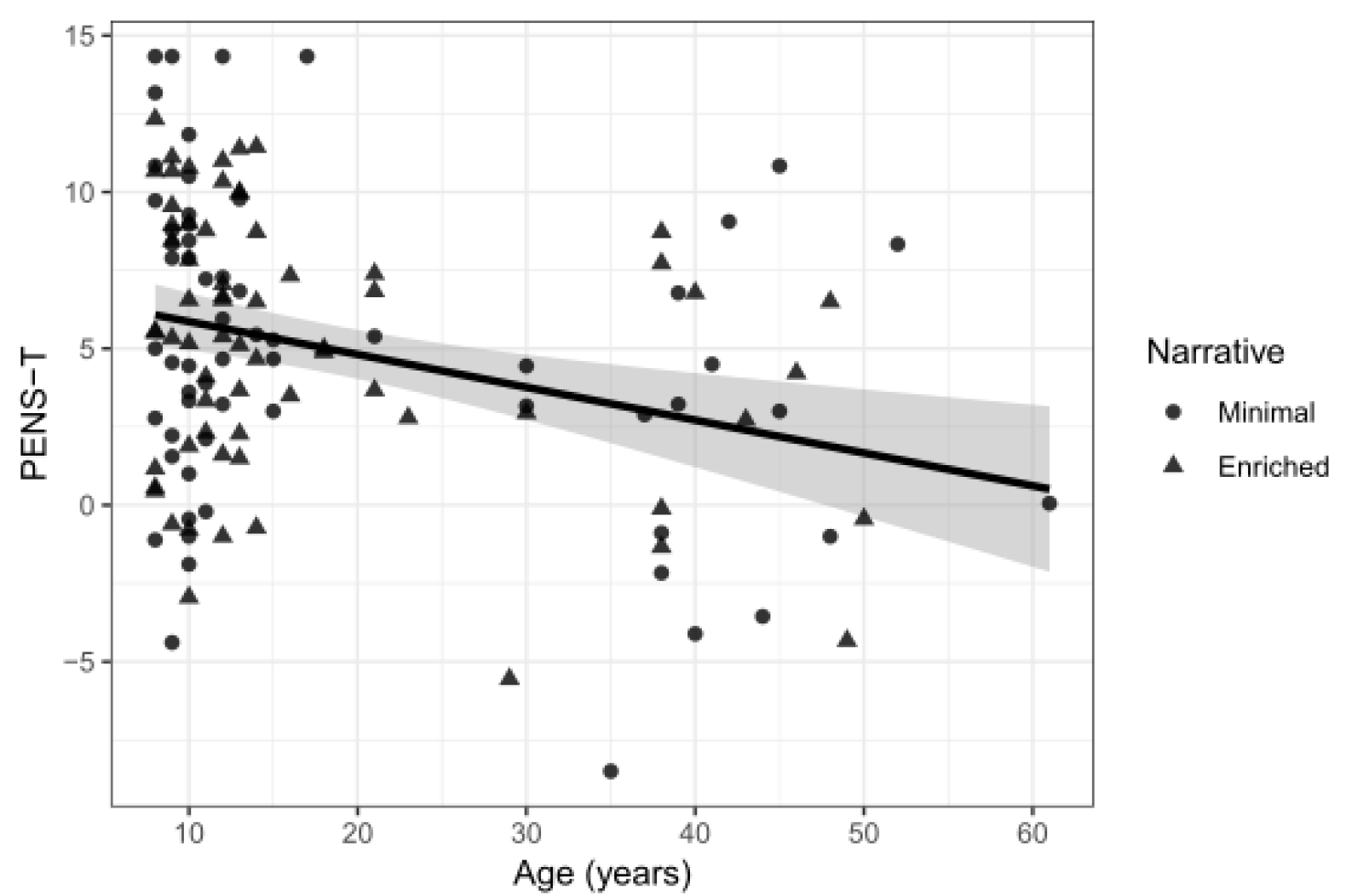
Scatterplot indicating negative relationship between age and PENS-T for the minimal (circles) and enriched (triangles) narrative groups. Shaded area corresponds to 95% confidence intervals for the linear trend.

In addition, age was negatively correlated with SIP scores, indicating higher presence for young participants (ρ(126) = −.26, *p* = .003).

Age was not correlated with the following measures: FMS (ρ(126) = .11, *p* = .22), PENS-PI (ρ(126) = −.16, *p* = .08), PENS-A (ρ(126) = −.15, *p* = .08), PENS-R (ρ(126) = −.16, *p* = .08), SSQ-T (ρ(126) = .01, *p* = .91), SSQ-N (ρ(126) = .04, *p* = .69), SSQ-O (ρ(126) = −.05, *p* = .57), SSQ-D (ρ(126) = .04, *p* = .65).

##### Sex

With respect to the main outcome variables, male and female participants did not differ with respect to presence (PENS-PI: *W* = 2100.5, *p* = .29) or cybersickness (SSQ-T: *W* = 2081, *p* = .33). In addition, we found no other effect of sex on outcome variables: SIP (*W* = 2261, *p* = .058), FMS (*W* = 1897, *p* = .96), SSQ-N (*W* = 2035, *p* = .45), SSQ-D (*W* = 2075.5, *p* = .33), SSQ-O (*W* = 2193, *p* = .12), PENS-T (*W* = 1775, *p* = .58), PENS-R (*W* = 1707.5, *p* = .37), PENS-A (*W* = 1873.5, *p* = .95), PENS-IC (*W* = 1619, *p* = .18), PENS-C (*W* = 1697, *p* = .35).

##### Cybersickness Profile

As in Experiment 1, we found that disorientation was the highest subscale score for the SSQ, followed by the nausea and oculomotor subscales (Figure 19).

**Figure 19.**
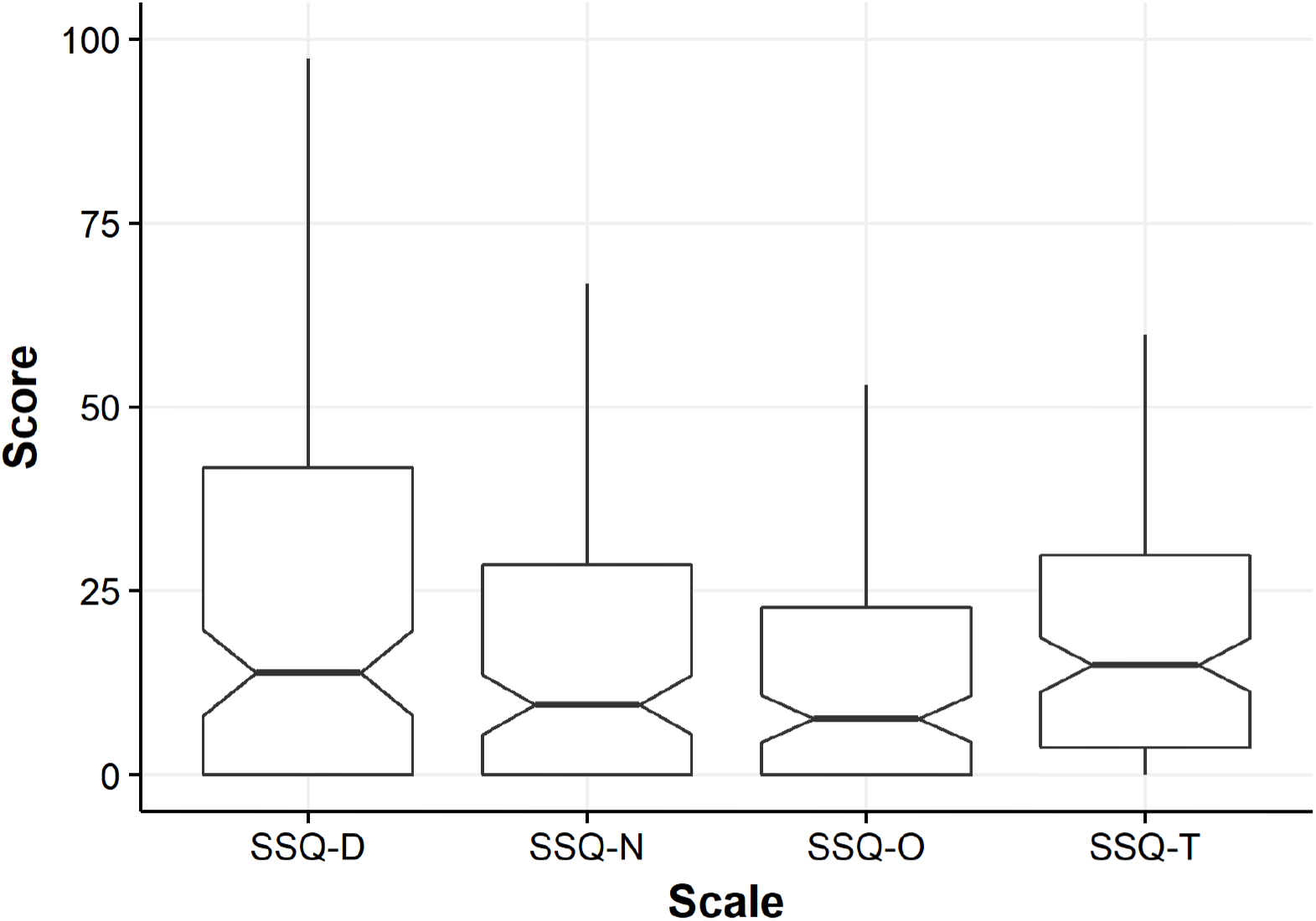
Notched Tukey boxplots showing cybersickness across subscales and total scores of the SSQ. Bars are IQRs. D = Disorientation. N = Nausea. O = Oculomotor, T = Total.

## General Discussion

### A-priori Hypotheses

Here we examined the relationship between presence and cybersickness in immersive VR, and assessed the potential effect of a narrative manipulation on these factors. Experiment 1 was conducted in a controlled laboratory setting. We anticipated a negative relationship between our primary measures of presence (PENS-PI) and cybersickness (SSQ-T), and we expected that an enriched narrative would produce lower cybersickness and higher presence. The results of Experiment 1 confirmed an inverse association between presence and cybersickness, in line with previous research and our predictions. In addition, we observed a higher level of presence in participants who received the enriched narrative in Experiment 1. However, the difference in cybersickness between the narrative groups was not significant. Interestingly, the relationship between presence and cybersickness was negative for the enriched narrative group, but not for the minimal narrative group, where the correlation was null. In Experiment 2 we tested if these effects held in a heterogeneous convenience sample that we collected at a public museum. Here, we once more confirmed that individuals who indicated that they experienced higher presence also reported lower levels of cybersickness, and vice-versa. However, the narrative manipulation had no effect on presence or cybersickness in the publicly-collected convenience sample.

#### Presence and cybersickness are negatively correlated

Across Experiment 1 and 2, we observed significant negative correlations between presence and cybersickness. At the same time, we identified no significant relationship between the rapid measures of presence and cybersickness in either experiment. Figures 8 and 14 reveal that responses tended to cluster at high levels of the SIP and low levels of the FMS. The conflict between this finding and the significant correlation between the SSQ-T and PENS-PI we found in both experiments aptly reflects the mixed literature on presence and cybersickness, where several studies have identified no relationship between the two concepts (see Weech et al., 2019, for a review).

#### Effects of a narrative manipulation in VR

We observed an expected effect of an enriched narrative on presence in Experiment 1. Although narrative context affected presence in Experiment 1, it did not influence other factors significantly, including the likelihood that participants would successfully complete their given task in VR (also in Experiment 2). This result aligns with our prediction that enriched narrative encourages engagement with the simulated environment as if it were real, directing attention away from artefacts of the simulation (e.g., audio-visual fidelity, character realism) or their own physiological response to multisensory conflicts in VR. Although we expected that this enhanced level of presence in the enriched narrative group would reduce cybersickness through the same theorized conflict-attenuation mechanism, we did not find evidence of a main effect of narrative on cybersickness in the same sample of participants. However, we found an interaction between narrative and gaming experience in terms of cybersickness (see *Exploratory analyses: The effect of narrative on cybersickness depends on video gaming experience*).

Contrary to our predictions, the results indicated that a narrative intervention had no direct effect on cybersickness or presence in the convenience sample of Experiment 2. The failure to confirm our expected results (and the results of Experiment 1 with respect to presence) may be related to the large number of noise sources contained in this dataset. The public setting of Experiment 2 meant that the similarity of the experience across participants could not be maintained precisely (e.g., ambient sound levels varied over time). The wide heterogeneity of our sample, while providing a good test of the generalizability of the effects observed in Experiment 1, also contributed a source of noise. Although we collected data on a number of individual factors (e.g., age, video game experience), other unmeasured factors could have influenced our data. These include the time constraints of visitors to the museum, and the recent activities of participants immediately prior to conducting the experiment (e.g., some may have been dancing in another museum exhibit, while others may have been watching a video).

While it is possible that a decreased focus on the narrative for younger children could have reduced our ability to detect a main effect, we found no evidence that age interacted with narrative context on the main outcome measures. Therefore, it may be that the effect of narrative context observed in Experiment 1 depends on being administered in a controlled laboratory environment, although it is possible that other unmeasured factors that varied between the two experiments led to this discrepancy.

Although we found no main effect of narrative on presence or cybersickness in Experiment 2, we observed an interaction between narrative and video game experience on cybersickness. However, since we did not establish this interaction as an a-priori prediction, we discuss this finding below in the section on exploratory analyses (see *Exploratory analyses: Regular Gamers vs. Non-Gamers*).

### Exploratory Analyses

While the above analyses were determined a-priori, we also aimed to provide a detailed picture of the relationships between cybersickness, presence, narrative, and other individual differences. For this reason, we collected indices of video game experience, and demographic factors (age, sex).

#### The presence-cybersickness correlation depends on narrative

In both experiments we found that the negative correlation between presence and cybersickness was significant when measured across all participants. However, when the minimal and enriched narrative groups were analyzed separately, we observed a significant negative trend only for the enriched narrative group. This dependency was unexpected and an explanation does not readily emerge on the basis of the current dataset.

The negative relationship between presence and cybersickness has been attributed to individual differences in the processing of multisensory conflicts: Individuals with a high degree of sensitivity to conflict tend to experience low presence and high cybersickness, and vice versa (Weech et al., 2019). An enriched narrative may enhance the likelihood that users feel immersed in an environment, thus diminishing the likelihood that the user will attend to features of the environment such as graphical fidelity and physics realism (rather, they are engaged with the story, characters, etc.). For individuals with low conflict sensitivity, enriched narrative is expected to enhance presence and reduce cybersickness due to the decreased focus on features external to the simulated environment. On the other hand, this immersiveness may exacerbate the sensory conflicts for individuals who are highly sensitive to conflicts, because the conflicts are less readily attributed as being artifacts of the simulation. This increase in cybersickness may subsequently detract from the feeling of presence as attention becomes directed inwards.

This proposed mechanism through which narrative context modulates the presence cybersickness relationship (for high conflict sensitive users: enriched narrative increases presence, leading to increased cybersickness, which subsequently reduces presence) could only be confirmed by collecting cybersickness and presence measured throughout a session of VR (we did not collect those data here). In addition, we recommend that individual sensitivities for sensory conflict should be measured prior to VR exposure and compared to presence (expected to be high for low-conflict sensitive individuals) and cybersickness (expected to be low for low-conflict sensitive individuals) measures.

#### The effect of narrative on cybersickness depends on video gaming experience

In both experiments we found that enriched narrative was associated with increased presence, but that the reductive effect of narrative on cybersickness depended on video gaming experience. Non-gamers reported higher cybersickness when receiving the minimal narrative, while regular gamers tended to exhibit similar levels of cybersickness regardless of the narrative context. This effect was strongest in Experiment 2, where we included a large sample of participants, but we also obtained a significant narrative-gaming experience interaction in Experiment 1.

The fact that the reductive effect of narrative on cybersickness only occurred for non-gamers suggests that gamers were already at relatively floor-levels of cybersickness, whereas non-gamers—who experienced greater cybersickness overall—benefited from an enriched narrative context. This finding has important implications for future interventions using narrative context to combat cybersickness and presence: While non-gamers constitute appropriate targets for such an approach, regular gamers might not benefit from the intervention.

#### Additional measures of presence and cybersickness

We found limited evidence in support of the idea that the rapid measures of presence and cybersickness were negatively correlated. Indeed, our results indicated that the vast majority of responses clustered at high levels of presence and low levels of cybersickness for these brief verbal reports. This pattern of results was unlike the trend we observed for the comprehensive measures of presence and cybersickness, where many participants had either low-presence and high-cybersickness, or high-presence and low-cybersickness. This factor may explain some of the discrepant findings from studies of the association between presence and cybersickness (Weech et al., 2019): Measurement techniques strongly modulate the type of relationship observed. We reason that the single item measures of presence and cybersickness are likely to encompass other experiential variables such as enjoyment (likely associated with high presence and low cybersickness), whereas the comprehensive measures of our outcome variables allow characterization of multiple sub-components of each (e.g., ‘oculomotor’ vs. ‘nausea’ types of cybersickness). We observed the strongest correlation between the FMS and the nausea subscale of the SSQ, although the variance explained was low-to-moderate (Exp. 1: 8%; Exp. 2: 47%). Similarly, the variance in SIP scores explained by the PENS-PI scale was low (Exp. 1: 24%; Exp. 2: 21%), reflecting that the two measures are not redundant. Our results reinforce the idea that it is highly desirable to obtain multiple outcome measures in studies of presence/cybersickness.

#### Regular gamers vs. Non-gamers

In Experiment 2 we observed that individuals who reported that they played games regularly indicated a higher intuitiveness of controls for the VR application, as well as higher competence in the game, likely reflecting a broad set of gaming skills developed across repeated sessions of video game play. Regular gamers were also less likely to report nausea and oculomotor discomfort symptoms, although presence was unaffected by previous gaming experience.

These results support previous evidence of an inverse correlation between video game experience and cybersickness levels (Knight & Arns, 2006). Note that other studies have failed to document any relationship between these factors (Gamito et al., 2010; 2008). Similarly, our data reflect the results of previous studies that found no evidence for a link between video game experience and presence (Alsina-Jurnet & Gutierrez-Maldonado, 2010; Ling et al., 2013; Schuemie et al., 2005), thus failing to support research showing that training with video games leads to enhanced levels of presence (Gamito et al., 2010).

#### Goal completion

We observed that participants who successfully achieved the goal of the task during VR exposure reported greater competence scores across Experiments 1 and 2, and scored higher on the ‘relatedness’ (connectedness to other in-game characters) and ‘total’ scores of the PENS scale in Experiment 2. While the difference in competence likely emerged due to the completion of the task, the effect observed for relatedness scores may reflect the fact that in-game characters engaged the user in conversation following completion of the task, generating a greater sense of character realism.

Surprisingly, we found that individuals in Experiment 1 who successfully completed the task also scored higher on the disorientation and oculomotor discomfort subscales of the SSQ. While we are limited to speculation about the behavior of participants (we did not collect quantitative data regarding movement in the environment), one explanation for this effect may be that individuals who completed the task did so because they explored the environment moreso and/or faster than other participants. This would have led to a higher chance of locating and activating the correct terminal, but also, an increased exposure to multisensory conflicts produced by passive self-translation in VR, generating cybersickness (Weech et al., 2019). An additional study that examines participants’ movement profiles would help to assess this hypothesis.

#### Effects of participant age

The sample of Experiment 2 demonstrated a wide age range, and our results supported an effect of age on game experience and rapid self-reports of presence. First, we identified that younger participants reported higher intuitiveness of controls and competence than older adults, and that these subscale differences led to a more positive overall score on the PENS game experience scale.

In previous research involving a large age-diverse sample (Vella et al., 2015; *N* = 297, 12-52 years, *M* age = 25.6, *SD* age = 8.0) age was significantly negatively correlated with several subscales of the PENS (presence-immersion, competence, and relatedness; note, the intuitive controls subscale was not calculated), and our data partially support this finding. While the previous study examined memory-recall of the participants’ most recently played game, our results confirm a relationship between game experience factors and age in a large sample for a standardized video game application.

Second, our results show that younger participants reported higher presence than older adults, but only for the single item measurement of presence taken immediately after immersion in VR. No age effect was observed for the more comprehensive PENS subscale of presence.

Evidence of enhanced susceptibility to presence in young individuals has been provided by previous research (Baumgartner et al., 2008; Schaik et al., 2004). In an fMRI study, an increased susceptibility to presence for children was supported by heightened bilateral amygdala and insula activation in children (*M* age = 8.7, *SD* age = 1.6, range = 6 to 11) than adults (*M* age = 26.3, *SD* age = 4.1, range = 21 to 43), indicating greater affective processing of an immersive virtual environment. At the same time, decreased prefrontal cortex activity in children was taken as evidence that, unlike adults, children did not tend to critically evaluate the virtual environment fidelity, which is a process that can cause breaks in presence. Baumgartner and colleagues (2008) also proposed that children are likely less able to suppress powerful multisensory stimuli that facilitate presence illusions. While our comprehensive measure of presence did not capture this previously documented relationship between age and presence, it is possible that some additional noise was added to this measure by the difficulty of very young children (~8 years of age) with focusing on completion of the questionnaire.

#### No effects of participant sex

Here, we found no evidence supporting a sex difference with respect to cybersickness, presence, or any other outcome measures that we gathered. While a variety of studies have found that levels of presence are higher either for women (Gamito et al., 2008) or for men (Felnhofer et al., 2012; Lachlan & Krcmar, 2011; Nicovich et al., 2005; Slater et al., 1998), our results agree with other research that found no effect of sex on presence ratings (De Leo et al., 2014). At the same time, while women tend to demonstrate higher cybersickness magnitude than men (De Leo et al., 2014; Häkkinen et al., 2002; Jaeger & Mourant, 2011; Park et al., 2006), the data obtained in our study agree with the work of others who identified no sex differences in cybersickness (Knight & Arns, 2006; Gamito et al., 2008; Ling et al., 2013).

#### Cybersickness profile

With respect to cybersickness data from the SSQ obtained in the samples of both Experiment 1 and 2, we identified a similar pattern of subscale scores as previous studies (e.g., Stanney et al., 1997; Weech et al., 2018b). The results of Stanney and colleagues (1997) revealed that cybersickness differs quantitatively from other forms of motion sickness (e.g., simulator sickness in motion-base simulators): The disorientation subscale score was higher than the others in cybersickness, whereas simulator sickness is predominantly characterized by oculomotor discomfort, with nausea being the intermediate magnitude for both conditions. The cybersickness data from the current study mirror this established finding.

#### Limitations of the study

Despite our large sample in Experiment 2, the analyses of this study were restricted with respect to statistical power. The number of participants who reported that they did not regularly play video games was low compared to our overall sample size, and our between-groups design for the narrative manipulation also limited our power, particularly in this group. This limitation, in addition to the fact that our exploratory analyses were not corrected for multiple comparisons, leads to important qualifiers about the strength of our conclusions on the basis of the current data. However, our sample size was very large for empirical research in this field (*N* = 128 in Experiment 2, 170 participants overall), as evidenced by a recent review of the literature (Weech et al., 2019). This fact, when coupled with the conservative non-parametric tests conducted for pairwise comparisons and correlational analyses, leads us to conclude that our findings provide a robust foundation for future confirmatory research that aims to further study the interaction of top-down and individual factors in the experience of virtual reality applications.

Another limitation involves the fact that participants were free to explore the VR environment as they saw fit, rather than being guided in a linear path. Although we selected a portion of the game that occurred in a constricted environmental space that participants were locked within, informal observations of participant behavior suggested that some individuals took the choice to remain relatively still in the environment, while others were freely exploring their environment. These individual differences generated high variance in the degree to which participants were exposed to the multisensory conflicts that have been strongly implicated in both presence, which is diminished by conflicts, and cybersickness, which tends to increase with large multisensory conflicts. In a future replication, it would be desirable to collect data using a VR experience that is more closely matched across participants, in addition to collecting data about the movement of participants and their virtual avatars. However, a collaborative effort between game developers and experimentalists will be required in order to maintain the high graphical fidelity with respect to the characters and environment that was achieved with the VR application used here.

## Conclusion

We assessed the association between presence and cybersickness, key factors of the VR experience, as well as their dependence on other factors such as narrative context, game experience, and demographic factors. Results of a laboratory experiment revealed a negative correlation between questionnaire measures of presence and cybersickness, but that the relationship was only significant for an enriched narrative context. Increased presence was reported after the delivery of an enriched narrative context, although enriched narrative only reduced cybersickness for non-gamers. In a second experiment involving a convenience sample from a museum, we supported our previous evidence of a negative correlation between presence and cybersickness for the enriched narrative group, and an interaction between narrative and gaming experience on cybersickness scores. The findings are complementary to recent suggestions that presence and cybersickness can be strongly modulated by individual factors that tend not to be measured in empirical studies in this field.

## Acknowledgements

This research was supported by grants from Oculus Research and the Natural Sciences and Engineering Research Council of Canada (NSERC) [Grant no. RGPIN-05435-2014] awarded to MB-C. The funding sources had no input in the preparation of the manuscript.

## Declarations of interests

Declarations of interest: none

## References

Abeele, V.V., Nacke, L.E., Mekler, E.D., Johnson, D., 2016. Design and preliminary validation of the Player Experience Inventory. CHI PLAY Companion’16, 335–341

Alsina-Jurnet, I., Gutiérrez-Maldonado, J., 2010. Influence of personality and individual abilities on the sense of presence experienced in anxiety triggering virtual environments. Int. J. Hum. Comput. Stud. 68(10), 788–801.

Bahit, M., Wibirama, S., Nugroho, H.A., Wijayanto, T., Winadi, M.N., 2016, October. Investigation of visual attention in day-night driving simulator during cybersickness occurrence. In Information Technology and Electrical Engineering (ICITEE), 2016 8th International Conference on (pp. 1–4). IEEE.

Bailey, J.H., Witmer, B.G., 1994, October. Learning and transfer of spatial knowledge in a virtual environment. In Proceedings of the Human Factors and Ergonomics Society Annual Meeting (Vol. 38, No. 18, pp. 1158–1162). Sage CA: Los Angeles, CA: SAGE Publications.

Baños, R.M., Botella, C., Alcañiz, M., Liaño, V., Guerrero, B., Rey, B., 2004. Immersion and emotion: their impact on the sense of presence. Cyberpsychol. Behav. 7(6), 734–741.

Baumgartner, T., Speck, D., Wettstein, D., Masnari, O., Beeli, G., Jäncke, L., 2008. Feeling present in arousing virtual reality worlds: prefrontal brain regions differentially orchestrate presence experience in adults and children. Front. Hum. Neurosci. 2, 8.

Bouchard, S., Robillard, G., St-Jacques, J., Dumoulin, S., Patry, M.J., Renaud, P., 2004, October. Reliability and validity of a single-item measure of presence in VR. In Proceedings of The 3rd IEEE International Workshop on Haptic, Audio and Visual Environments and Their Applications, HAVE 2004. (pp. 59–61). IEEE.

Bouchard, S., St-Jacques, J., Robillard, G., Renaud, P., 2008. Anxiety increases the feeling of presence in virtual reality. Pres. Teleop. Virtual Environ. 17(4), 376–391.

Brockmyer, J.H., Fox, C.M., Curtiss, K.A., McBroom, E., Burkhart, K.M., Pidruzny, J.N., 2009. The development of the Game Engagement Questionnaire: A measure of engagement in video game-playing. J. Exp. Soc. Psychol. 45(4), 624–634.

Busscher, B., De Vliegher, D., Ling, Y., & Brinkman, W. P., 2011. Physiological measures and self-report to evaluate neutral virtual reality worlds. J. Cyber. Ther. Rehabil. 4(1), 1–18.

Chelen, W.E., Kabrisky, M., Rogers, S. K., 1993. Spectral analysis of the electroencephalographic response to motion sickness. Aviat. Space Environ. Med. 64(1), 24.

Cobb, S.V., Nichols, S., Ramsey, A., Wilson, J.R., 1999. Virtual reality-induced symptoms and effects (VRISE). Pres. Teleop. Virtual Environ. 8(2), 169–186.

Cooper, N., Milella, F., Cant, I., Pinto, C., White, M.D., Meyer, G.F., 2016. The effects of multisensory cues on the sense of presence and task performance in a virtual reality environment. In Perception (Vol. 45, pp. 332–333). Sage Publications LTD, London, England.

Davis, S., Nesbitt, K., Nalivaiko, E., 2015, January. Comparing the onset of cybersickness using the Oculus Rift and two virtual roller coasters. In Proceedings of the 11th Australasian Conference on Interactive Entertainment (IE 2015) (Vol. 27, p. 30).

De Leo, G., Diggs, L.A., Radici, E., Mastaglio, T.W., 2014. Measuring sense of presence and user characteristics to predict effective training in an online simulated virtual environment. Simul. Healthc. 9(1), 1–6.

Felnhofer, A., Kothgassner, O.D., Beutl, L., Hlavacs, H., Kryspin-Exner, I., 2012. Is virtual reality made for men only? Exploring gender differences in the sense of presence. In Proceedings of the International Society on Presence Research, 103–112.

Friedman, R.S., Förster, J., 2010. Implicit affective cues and attentional tuning: an integrative review. Psychol. Bull. 136(5), 875.

Gamito, P., Oliveira, J., Morais, D., Baptista, A., Santos, N., Soares, F., Saraiva, T., Rosa, P., 2010. Training presence: The importance of virtual reality experience on the “sense of being there”. Ann. Rev. Cyberther. Telemed. 2010, 128–133.

Gamito, P., Oliveira, J., Santos, P., Morais, D., Saraiva, T., Pombal, M., Mota, B., 2008. Presence, immersion and cybersickness assessment through a test anxiety virtual environment. Ann. Rev. Cyberther. Telemed. 2008, 83–90.

Guttentag, D.A., 2010. Virtual reality: Applications and implications for tourism. Tourism Manage. 31(5), 637–651.

Häkkinen, J., Vuori, T., Paakka, M., 2002, October. Postural stability and sickness symptoms after HMD use. In IEEE International Conference on Systems, Man and Cybernetics (Vol. 1, pp. 147–152). IEEE.

Jaeger, B.K., Mourant, R.R., 2001, October. Comparison of simulator sickness using static and dynamic walking simulators. In Proceedings of the Human Factors and Ergonomics Society Annual Meeting (Vol. 45, No. 27, pp.1896–1900). Sage CA: Los Angeles, CA: SAGE Publications.

Jin, W., Choo, A., Gromala, D., Shaw, C., Squire, P., 2016, April. A Virtual Reality Game for Chronic Pain Management: A Randomized, Controlled Clinical Study. In MMVR (pp. 154–160).

Kennedy, R.S., Lane, N.E., Berbaum, K.S., & Lilienthal, M.G., 1993. Simulator sickness questionnaire: An enhanced method for quantifying simulator sickness. Int. J. Aviat. Psychol. 3(3), 203–220.

Keshavarz, B., Hecht, H., 2011. Validating an efficient method to quantify motion sickness. Hum. Factors 53(4), 415–426.

Knight, M.M., Arns, L.L., 2006, July. The relationship among age and other factors on incidence of cybersickness in immersive environment users. In ACM SIGGRAPH 2006 Research Posters (p.196). ACM.

Lachlan, K., Krcmar, M., 2011. Experiencing presence in video games: The role of presence tendencies, game experience, gender, and time spent in play. Commun. Res. Rep. 28(1), 27–31.

LaViola Jr, J.J., 2000. A discussion of cybersickness in virtual environments. ACM SIGCHI Bulletin, 32(1), 47–56.

Lin, J.H.T., 2017. Fear in virtual reality (VR): Fear elements, coping reactions, immediate and next-day fright responses toward a survival horror zombie virtual reality game. Comput. Hum. Behav. 72, 350–361.

Lin, J.W., Duh, H.B.L., Parker, D.E., Abi-Rached, H., Furness, T.A., 2002. Effects of field of view on presence, enjoyment, memory, and simulator sickness in a virtual environment. In IEEE Proceedings of Virtual Reality, 2002 (pp.164–171). IEEE.

Ling, Y., Nefs, H.T., Brinkman, W.P., Qu, C., Heynderickx, I., 2013. The relationship between individual characteristics and experienced presence. Comput. Hum. Behav. 29(4), 1519–1530.

Liu, C.L., Uang, S.T., 2011. Effects of presence on causing cybersickness in the elderly within a 3D virtual store. In International Conference on Human-Computer Interaction, July 2011 (pp.490–499). Springer: Cham, Switzerland.

McCauley, M.E., Sharkey, T.J., 1992. Cybersickness: Perception of self-motion in virtual environments. Pres. Teleop. Virtual Environ. 1(3), 311–318.

Melo, M., Sampaio, S., Barbosa, L., Vasconcelos-Raposo, J., Bessa, M., 2016, November. The impact of different exposure times to 360° video experience on the sense of presence. In Computação Gráfica e Interação (EPCGI), 2016 23° Encontro Português de (pp. 1–5). IEEE.

Melo, M., Vasconcelos-Raposo, J., Bessa, M., 2018. Presence and cybersickness in immersive content: Effects of content type, exposure time and gender. Comput. Graph. 71, 159–165.

Miller, A.D., Rowley, H.A., Roberts, T.P., Kucharczyk, J., 1996. Human Cortical Activity during Vestibular‐and Drug‐Induced Nausea Detected Using MSIa. Ann. N. Y. Acad. Sci. 781(1), 670–672.

Money, K.E., Wood, J.D., 1970, January. Neural mechanisms underlying the symptomatology of motion sickness. In Fourth Symposium on the role of the vestibular organs in space exploration (pp. 69–82). NASA, Houston.

Nichols, S., Haldane, C., Wilson, J.R., 2000. Measurement of presence and its consequences in virtual environments. Int. J. Hum. Comput. Stud. 52(3), 471–491.

Nicovich, S.G., Boller, G.W., Corwell, T.B., 2005. Experienced presence within computer-mediated communications: initial explorations on the effects of gender with respect to empathy and immersion. J. Comput. Mediat. Comm. 10(2).

Nielsen, L.T., Møller, M.B., Hartmeyer, S.D., Ljung, T., Nilsson, N.C., Nordahl, R., Serafin, S., 2016, November. Missing the point: an exploration of how to guide users’ attention during cinematic virtual reality. In Proceedings of the 22nd ACM Conference on Virtual Reality Software and Technology (pp. 229–232). ACM.

Oman, C.M., 1990. Motion sickness: a synthesis and evaluation of the sensory conflict theory. Can. J. Physiol. Pharmacol. 68(2), 294–303.

Palmisano, S., Mursic, R., Kim, J., 2017. Vection and cybersickness generated by head-and-display motion in the Oculus Rift. Displays 46, 1–8.

Park, G.D., Allen, R.W., Fiorentino, D., Rosenthal, T.J., Cook, M.L., 2006. Simulator sickness scores according to symptom susceptibility, age, and gender for an older driver assessment study. In Proceedings of the human factors and ergonomics society annual meeting, October 2006 (Vol. 50, No. 26, pp.2702–2706). Sage CA: Los Angeles, CA, USA.

Reason, J.T., 1978. Motion sickness adaptation: a neural mismatch model. J. R. Soc. Med. 71(11), 819–829.

Rebenitsch, L., Owen, C., 2016. Review on cybersickness in applications and visual displays. Virtual Real. 20(2), 101–125.

Reschke, M.F., Bloomberg, J.J., Harm, D.L., Paloski, W.H., Layne, C., McDonald, V., 1998. Posture, locomotion, spatial orientation, and motion sickness as a function of space flight. Brain Res. Rev. 28(1-2), 102–117.

Ryan, R.M., Rigby, C.S., Przybylski, A., 2006. The motivational pull of video games: A self-determination theory approach. Motiv. Emot. 30(4), 344–360.

Sanchez-Vives, M.V., Slater, M., 2005. From presence to consciousness through virtual reality. Nat. Rev. Neurosci. 6(4), 332.

Schuemie, M.J., Abel, B., van der Mast, C.A.P.G., Krijn, M., Emmelkamp, P.M.G., 2005. The effect of locomotion technique on presence, fear and usability in a virtual environment. In EUROMEDIA (Vol. 2005, p.11).

Schuemie, M.J., Van Der Straaten, P., Krijn, M., Van Der Mast, C.A., 2001. Research on presence in virtual reality: A survey. Cyberpsychol. Behav. 4(2), 183–201.

Schaik, P.V., Turnbull, T., Wersch, A.V., Drummond, S., 2004. Presence within a mixed reality environment. Cyberpsychol. Behav. 7(5), 540–552.

Slater, M., Steed, A., McCarthy, J., Maringelli, F., 1998. The influence of body movement on subjective presence in virtual environments. Hum. Factors 40, 469–477.

Smolentsev, A., Cornick, J.E., Blascovich, J., 2017. Using a preamble to increase presence in digital virtual environments. Virtual Real. 21(3), 153–164.

Stanney, K.M., Hash, P., 1998. Locus of user-initiated control in virtual environments: Influences on cybersickness. Presence 7(5), 447–459.

Stanney, K.M., Kennedy, R.S., Drexler, J.M., 1997, October. Cybersickness is not simulator sickness. In Proceedings of the Human Factors and Ergonomics Society annual meeting (Vol. 41, No. 2, pp. 1138–1142). Sage CA: Los Angeles, CA: SAGE Publications.

Stanney, K.M., Kennedy, R.S., Drexler, J.M., Harm, D.L., 1999. Motion sickness and proprioceptive aftereffects following virtual environment exposure. Appl. Ergon. 30(1), 27–38.

Steinicke, F., Bruder, G., Hinrichs, K., Steed, A., 2010. Gradual transitions and their effects on presence and distance estimation. Comput. Graph. 34(1), 26–33.

Steinicke, F., Bruder, G., Hinrichs, K., Steed, A., Gerlach, A.L., 2009, March. Does a gradual transition to the virtual world increase presence? In Virtual Reality Conference, 2009. VR 2009. IEEE (pp. 203–210). IEEE.

Steuer, J., 1992. Defining virtual reality: Dimensions determining telepresence. J. Comm. 42(4), 73–93.

Stróżak, P., Francuz, P., Lewkowicz, R., Augustynowicz, P., Fudali-Czyż, A., Bałaj, B., Truszczyński, O., 2018. Selective Attention and Working Memory Under Spatial Disorientation in a Flight Simulator. Int. J. Aerospace Psychol. 28, 31–45.

Takahashi, M., Ogata, M., Miura, M., 1997. The significance of motion sickness in the vestibular system. J. Vestib. Res. 7(2-3), 179–187.

Usoh, M., Arthur, K., Whitton, M.C., Bastos, R., Steed, A., Slater, M., Brooks Jr, F.P., 1999. Walking> walking-in-place> flying, in virtual environments. In Proceedings of the 26th Annual Conference on Computer Graphics and Interactive Techniques, July 1999 (pp.359–364). ACM Press/Addison-Wesley Publishing Company.

van Emmerik, M.L., de Vries, S.C., Bos, J.E., 2011. Internal and external fields of view affect cybersickness. Displays 32(4), 169–174.

Vanden Abeele, V., Nacke, L.E., Mekler, E.D., Johnson, D., 2016, October. Design and preliminary validation of the player experience inventory. In Proceedings of the 2016 Annual Symposium on Computer-Human Interaction in Play Companion Extended Abstracts (pp. 335–341). ACM.

Vella, K., Johnson, D., Hides, L., 2015, December. Indicators of wellbeing in recreational video game players. In Proceedings of the Annual Meeting of the Australian Special Interest Group for Computer Human Interaction (pp. 613–617). ACM.

Weech, S., Kenny, S., & Barnett-Cowan, M., 2019. Presence and cybersickness in virtual reality are negatively related: A review. Front. Psychol. 10(158), 1–19.

Weech, S., Moon, J. & Troje, N.F., 2018a. Influence of bone-conducted vibration on simulator sickness in virtual reality. PloS One, 13(3), p.e0194137.

Weech, S., Varghese, J.P., & Barnett-Cowan, M., 2018b. Estimating the sensorimotor components of cybersickness. J. Neurophysiol. 120(5), 2201–2217.

Wilson, J.R., Nichols, S., Haldane, C., 1997. Presence and side effects: Complementary or contradictory? In Proc. of the HCI’97, San Francisco (pp. 889–892).

Witmer, B.G., Singer, M.J., 1998. Measuring presence in virtual environments: A presence questionnaire. Presence 7(3), 225–240.

